# Integration of genetic fine-mapping and multi-omics data reveals candidate effector genes for hypertension

**DOI:** 10.1101/2023.01.26.525702

**Authors:** Stefan van Duijvenboden, Julia Ramírez, William J. Young, Kaya J. Olczak, Farah Ahmed, Mohammed J.A.Y. Alhammadi, International Consortium of Blood Pressure, Christopher G. Bell, Andrew P. Morris, Patricia B. Munroe

## Abstract

Genome-wide association studies of blood pressure (BP) have identified >1000 loci but the effector genes and biological pathways at these loci are mostly unknown. Using published meta-analysis summary statistics, we conducted annotation-informed fine-mapping incorporating tissue-specific chromatin segmentation to identify causal variants and candidate effector genes for systolic BP, diastolic BP, and pulse pressure. We observed 532 distinct signals associated with ≥2 BP traits and 84 with all three. For >20% of signals, a single variant accounted for >75% posterior probability, 65 were missense variants in known (*SLC39A8, ADRB2, DBH*) and previously unreported BP candidate genes (*NRIP1, MMP14*). In disease-relevant tissues, we colocalized >80 and >400 distinct signals for each BP trait with *cis*-eQTLs, and regulatory regions from promoter capture Hi-C, respectively. Integrating mouse, human disorder, tissue expression data and literature review, we provide consolidated evidence for 394 BP candidate genes for future functional validation and identifies several new drug targets.

## Introduction

Elevated blood pressure (BP) affects over 1 billion people and is one of the most important risk-factors for cardiovascular disease (CVD), leading to significant mortality and morbidity worldwide^1^. It is estimated to cause more than 10 million deaths per year^2^. Genome-wide association studies (GWAS), bespoke targeted arrays (Cardio Metabochip) and Exome-array wide association studies (EAWAS) have identified over 1,000 BP-associated loci for this heritable polygenic trait^3–8^. However, for most loci, the effector genes and relevant biological processes through which BP associations are mediated have yet to be characterised. Here, we use published GWAS meta-analysis results (n >757,000) of systolic BP (SBP), diastolic BP (DBP) and pulse pressure (PP)^3^ to perform fine mapping of causal variants at BP loci. Through the integration of GWAS with tissue specific epigenomic annotations, and colocalisation of BP associations with expression quantitative loci (eQTLs) and Hi-C promoter interactions in relevant BP tissues, we identify high confidence effector genes and causal pathways, and assess their potential for drag target identification or repurposing opportunities

## Results

### Study overview and reporting of loci

We utilised previously reported meta-analyses of GWAS of BP traits in up to 757,601 individuals of European ancestry from the International Consortium of BP and UK Biobank (ICBP+UKBB)^3^. Each contributing GWAS had been imputed up to reference panels from the 1000 Genomes Project^9,10^ and/or Haplotype Reference Consortium^11^. After quality control, meta-analysis association summary statistics for SBP, DBP and PP were reported for up to 7,160,657 single nucleotide variants (SNVs). An overview of the study design is provided in Supplementary Figure 1.

We considered a total of 606 genomic regions encompassing previously reported lead SNVs for SBP, DBP or PP that attained genome-wide significance (*P* < 5×10^−8^) for at least one BP trait (**Methods**, Supplementary Table 1). Through approximate cross-trait conditional analyses (**Methods**), we partitioned BP associations at the 606 genomic regions into a total of 1,850 distinct signals that were associated with at least one BP trait at genome-wide significance (Figure 1, Supplementary Table 2). Of these signals, 532 were associated with at least two BP traits (333 with SBP and DBP, 267 with SBP and PP, and 100 with DBP and PP), and 84 were associated with all three traits. The only discordancy in direction of effect was for 17 of the 100 signals shared across DBP and PP, where the DBP increasing allele was the PP decreasing allele).

**Figure 1.**
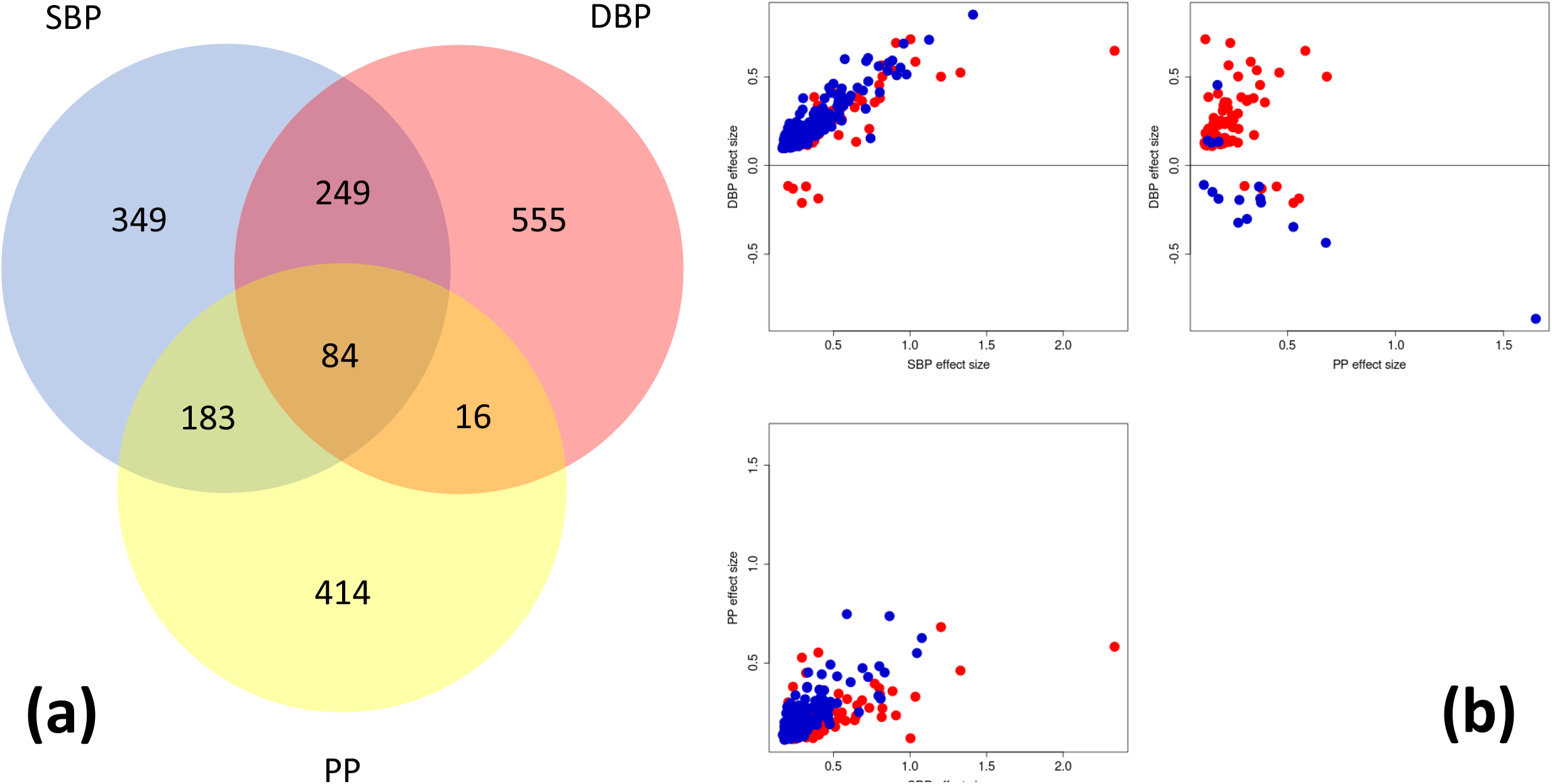
Overlap of 1,850 distinct signals attaining genome-wide significant evidence of association with SBP, DBP and PP in meta-analysis of BP GWAS in up to 757,601 individuals of European ancestry. (a) Venn diagram showing the number of signals shared across BP traits. Sharing of signals across traits is much more common between SBP and DBP or SBP and PP, with just 16 associations shared between only DBP and PP. (b) Comparison of allelic effect sizes on SBP, DBP and PP for the index SNV at the 532 distinct association signals that are shared across multiple BP traits. The effect has been aligned to the SBP or PP increasing allele for the signal. Blue points correspond to the 448 association signals that are shared across exactly two BP traits, whilst red points correspond to the 84 association signals that are shared across all three BP traits. When signals are shared between SBP and PP, the direction of effect of the index SNV on the traits is always concordant.

The cross-trait approximate conditional analyses revealed several genomic regions with complex patterns of associations with SBP, DBP and PP. For six genomic regions, more than 20 distinct signals of association were observed for at least one BP trait. The most complex associations were observed across: (i) a 6.4Mb region of chromosome 17, encompassing previously reported loci that include *PLCD3, GOSR2, HOXB7, ZNF652*, and *PHB* (locus ID 576, 37 distinct signals); (ii) a 5.8Mb region of chromosome 10, encompassing previously reported loci that include *PAX2, CYP17A1, NT5C2*, and *OBFC1* (locus ID 403, 34 distinct signals); and (iii) the major histocompatibility complex region of chromosome 6 (5.7Mb, locus ID 251, 32 distinct signals) that encompasses previously reported loci that include *BAT2, BAT5*, and *HLA DQB1*.

### Fine-mapping and genomic annotation reveals high-confidence causal variants for BP traits

Previous studies have demonstrated that improved localisation of causal variants driving association signals for complex human traits can be attained by integrating GWAS data with genomic annotation^12^. By mapping SNVs to functional and regulatory annotations from GENCODE^13,14^ and the Roadmap Epigenomics Consortium^15^ (**Methods**), we observed significant joint enrichment (*P* < 0.0002, Bonferroni correction for 253 annotations) for BP associations mapping to protein coding exons and 3’ UTRs, enhancers in heart and adrenal gland, and promoters in adipose and heart (Supplementary Table 3, Supplementary Figure 2).

For each distinct signal, we then derived credible sets of variants that together account for 99% of the posterior probability (π) of driving the BP trait association under an annotation-informed prior model of causality in which SNVs mapping to the genomic annotations in the globally enriched signatures for SBP, DBP and PP are upweighted (**Methods**). The median 99% credible set size was 20 variants for SBP and DBP, and 22 for PP (Supplementary Table 4). For 208 (24%), 224 (24.8%) and 159 (22.9%) SBP, DBP and PP signals, respectively, a single SNV accounted for more than 75% of the posterior probability of driving the BP association under the annotation-informed prior, which we defined as “high-confidence” for causality (Figure 2, Supplementary Table 5).

**Figure 2.**
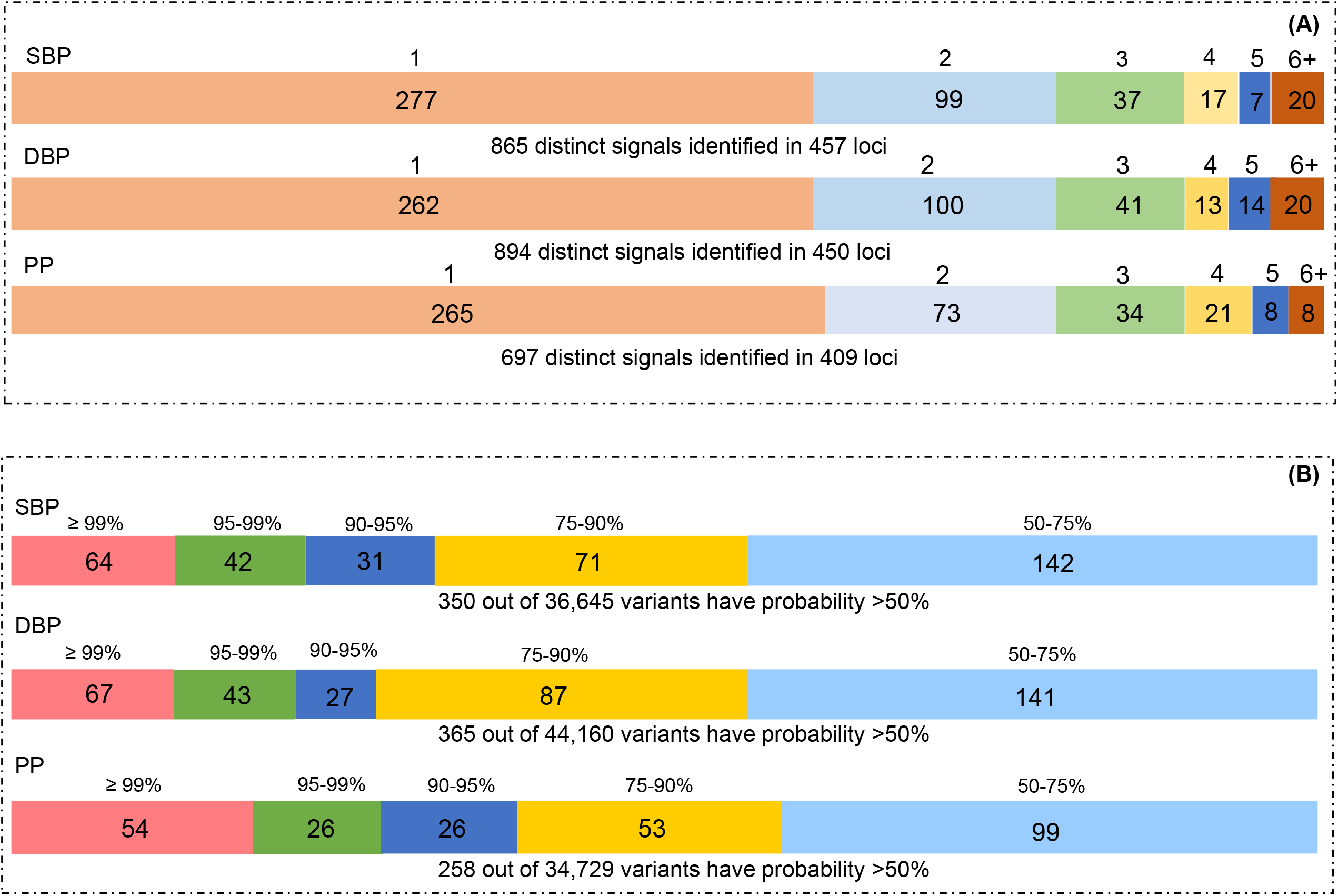
Distinct BP association signals. (a) Summary of distinct association signals for blood pressure traits. SBP: A single signal at 277 genomic regions and at least two at 180; DBP: A single signal at 262 genomic regions, and at least two at 188; PP: A single signal at 265 genomic regions, and at least two at 144. (b) Distribution of the posterior probability of causality of the variants in credible sets. SBP, systolic blood pressure; DBP, diastolic blood pressure; PP, pulse pressure.

### High-confidence SNVs are enriched for BP-related phenotypes

We used the Genomic Regions Enrichment of Annotations Tool (GREAT) v4.0.4^16^ (**Methods**) to explore the potential biological impact of all high confidence SNVs through their enrichment within trait-related genomic regions including *cis*-regulatory elements (CREs). We explored SNVs separately for the three BP traits and physiologically consistent enrichment results were identified for these location data for Gene Ontology Biological Processes (e.g., circulatory system processes, regulation of BP), Human Phenotype (e.g., abnormality systemic blood pressure, abnormality of vasculature), Mouse Phenotype and Knockout data (e.g., abnormal blood vessel morphology, increased systemic arterial blood pressure) (Supplementary Figure 3 and Supplementary Table 6).

### Missense variants implicate causal candidate genes

We identified 65 high-confidence missense variants for BP association signals (Supplementary Table 7). Among these, 20 were driving the same association signal across two BP traits, and one (*RGL3* p.Pro162His) was driving the same association signal across all three BP traits (Table 1). *RGL3* is not well characterised, but several missense variants in the gene have been previously identified in BP EAWAS^7^. In our study, three distinct association signals are driven by *HFE* missense variants (*HFE* p.His63Asp, *HFE* p. Ser65Cys and *HFE* p.Cys282Tyr). These variants are associated with predisposition to hereditary hemochromatosis, of which, portal hypertension and restrictive diastolic function are recognised phenotypes ^17^.

**Table 1.** **High-confidence missense variants for blood pressure association signals**

| Signal ID | Index SNV | Missense variant | Canonical transcript | Annotation | Chr | Position | Polyphen | SIFT | Trait | p-value | Posterior probability (%) |
| --- | --- | --- | --- | --- | --- | --- | --- | --- | --- | --- | --- |
| **1_6** | rs262695 | rs262695 | ENST00000545087.1 | *AL590822.1*<br>p.Cys78Arg | 1 | 2,144,788 | N/A | N/A | SBP | 9.4E-14 | 96.2 |
|  |  |  |  |  |  |  |  |  | PP | 1.6E-12 | 99.1 |
| **3_1** | rs11121483 | rs846111 | ENST00000377939.4 | *RNF207*<br>p.Gly603Ala | 1 | 6,279,370 | benign | tolerated (LC) | PP | 1.8E-10 | 83.5 |
| **39_1** | rs150816167 | rs61747728 | ENST00000367615.4 | *NPHS2*<br>p.Arg229Gln | 1 | 179,526,214 | possibly damaging | tolerated | DBP | 5.7E-11 | 88.1 |
| **52_1** | rs699 | rs699 | ENST00000366667.4 | *AGT*<br>p.Met268Thr | 1 | 230,845,794 | benign | tolerated | SBP | 5.6E-34 | 93.6 |
| **60_2** | rs2384061 | rs11676272 | ENST00000260600.5 | *ADCY3*<br>p.Ser107Pro | 2 | 25,141,538 | benign | tolerated | DBP | 5.4E-23 | 86.8 |
| **98_5** | rs10497529 | rs10497529 | ENST00000420890.2 | *CCDC141*<br>p.Ala141Val | 2 | 179,839,888 | probably damaging | deleterious | PP | 8.1E-18 | 89.0 |
| **108_1** | rs1047891 | rs1047891 | ENST00000430249.2 | *CPS1*<br>p.Thr1412Asn | 2 | 211,540,507 | benign | tolerated | SBP | 1.4E-14 | 98.5 |
|  |  |  |  |  |  |  |  |  | DBP | 8.2E-14 | 99.5 |
| **110_1** | rs1250259 | rs1250259 | ENST00000354785.4 | *FN1*<br>p.Gln15Leu | 2 | 216,300,482 | benign | tolerated | SBP | 7.5E-20 | 92.3 |
|  |  |  |  |  |  |  |  |  | PP | 8.6E-33 | 86.1 |
| **132_2** | rs74951356 | rs74951356 | ENST00000418109.1 | *LAMB2*<br>p.Ala1765Thr | 3 | 49,158,763 | benign | tolerated | DBP | 5.0E-09 | 95.8 |
| **135_1** | rs3772219 | rs3772219 | ENST00000338458.4 | *ARHGEF3*<br>p.Leu367Val | 3 | 56,771,251 | benign | tolerated | SBP | 3.1E-17 | 75.2 |
|  |  |  |  |  |  |  |  |  | DBP | 1.7E-24 | 81.1 |
| **158_3** | rs61762319 | rs61762319 | ENST00000460393.1 | *MME*<br>p.Met8Val | 3 | 154,801,978 | benign | deleterious | SBP | 1.5E-09 | 99.8 |
| **170_6** | rs2498323 | rs2498323 | ENST00000382774.3 | *HGFAC*<br>p.Arg644Gln | 4 | 3,451,109 | possibly damaging | tolerated | PP | 7.4E-13 | 100.0 |
| **174_1** | rs2302212 | rs2302212 | ENST00000514176.1 | *NCAPG*<br>p.Met231Thr | 4 | 17,818,885 | benign | deleterious | SBP | 4.1E-08 | 84.2 |
| **191_3** | rs13107325 | rs13107325 | ENST00000394833.2 | *SLC39A8*<br>p.Ala391Thr | 4 | 103,188,709 | possibly<br>damaging | tolerated | SBP | 4.2E-53 | 100.0 |
|  |  |  |  |  |  |  |  |  | DBP | 2.5E-90 | 100.0 |
| **221_1** | rs2307111 | rs2307111 | ENST00000428202.2 | *POC5*<br>p.His36Arg | 5 | 75,003,678 | benign | tolerated | DBP | 1.6E-22 | 97.6 |
| **237_6** | rs12189018 | rs1042713 | ENST00000305988.4 | *ADRB2*<br>p.Gly16Arg | 5 | 148,206,440 | benign | tolerated | DBP | 8.1E-09 | 81.7 |
| **237_7** | rs1800888 | rs1800888 | ENST00000305988.4 | *ADRB2*<br>p.Thr164Ile | 5 | 148,206,885 | benign | tolerated | DBP | 7.4E-13 | 100.0 |
| **249_4** | rs198851 | rs1799945 | ENST00000357618.5 | *HFE*<br>p.His63Asp | 6 | 26,091,179 | probably<br>damaging | tolerated | DBP | 5.7E-51 | 85.3 |
| **249_1** | rs1800730 | rs1800730 | ENST00000357618.5 | *HFE*<br>p.Ser65Cys | 6 | 26,091,185 | probably<br>damaging | deleterious | SBP | 2.0E-09 | 96.4 |
|  |  |  |  |  |  |  |  |  | DBP | 1.6E-18 | 100.0 |
| **249_3** | rs1800562 | rs1800562 | ENST00000357618.5 | *HFE*<br>p.Cys282Tyr | 6 | 26,093,141 | probably<br>damaging | deleterious | SBP | 2.0E-14 | 89.1 |
|  |  |  |  |  |  |  |  |  | DBP | 2.1E-37 | 96.4 |
| **251_7** | rs41543814 | rs41543814 | ENST00000376228.5 | *HLA-C*<br>p.Ala97Thr | 6 | 31,239,430 | benign | tolerated (LC) | DBP | 1.5E-19 | 100.0 |
| **251_10** | rs2844573 | rs2308655 | ENST00000412585.2 | *HLA-B*<br>p.Cys349Ser | 6 | 31,322,303 | benign | tolerated (LC) | PP | 1.4E-12 | 99.2 |
| **251_11** | rs1077393 | rs1052486 | ENST00000375964.6 | *BAG6*<br>p.Ser625Pro | 6 | 31,610,686 | benign | tolerated | DBP | 4.6E-24 | 86.1 |
| **252_1** | rs3176336 | rs2395655 | ENST00000448526.2 | *CDKN1A*<br>p.Asp28Gly | 6 | 36,645,696 | benign | tolerated (LC) | PP | 4.7E-13 | 98.5 |
| **255_1** | rs78648104 | rs78648104 | ENST00000008391.3 | *TFAP2D*<br>p.Phe74Leu | 6 | 50,683,009 | benign | tolerated | SBP | 2.4E-15 | 99.9 |
|  |  |  |  |  |  |  |  |  | DBP | 2.9E-20 | 100.0 |
| **272_1** | rs6919947 | rs6919947 | ENST00000368357.3 | *NCOA7*<br>p.Ser399Ala | 6 | 126,210,395 | benign | tolerated (LC) | SBP | 4.9E-17 | 100.0 |
|  |  |  |  |  |  |  |  |  | DBP | 2.2E-13 | 99.9 |
| **300_2** | rs2107732 | rs2107732 | ENST00000381112.3 | *CCM2*<br>p.Val74Ile | 7 | 45,077,978 | benign | tolerated | DBP | 3.3E-10 | 80.6 |
| **300_3** | rs2854746 | rs2854746 | ENST00000381083.4 | *IGFBP3*<br>p.Ala32Gly | 7 | 45,960,645 | benign | tolerated | DBP | 5.0E-11 | 97.9 |
| **313_2** | rs11556924 | rs11556924 | ENST00000358303.4 | *ZC3HC1*<br>p.Arg363His | 7 | 129,663,496 | probably<br>damaging | deleterious | DBP | 1.5E-26 | 98.3 |
| **313_4** | rs13222308 | rs3734928 | ENST00000335420.5 | *KLHDC10*<br>p.Ser2Leu | 7 | 129,710,488 | benign | deleterious<br>(LC) | SBP | 1.3E-10 | 90.0 |
| **353_10** | rs34591516 | rs34591516 | ENST00000377741.3 | *GPR20*<br>p.Gly313Ser | 8 | 142,367,087 | benign | tolerated | DBP | 1.3E-15 | 80.8 |
| **365_2** | rs76452347 | rs76452347 | ENST00000354323.2 | *HRCT1*<br>p.Arg63Trp | 9 | 35,906,471 | possibly<br>damaging | deleterious<br>(LC) | SBP | 7.1E-14 | 100.0 |
|  |  |  |  |  |  |  |  |  | DBP | 8.0E-23 | 100.0 |
| **372_3** | rs3739451 | rs3739451 | ENST00000401783.2 | *SVEP1*<br>p.Phe3161Ile | 9 | 113,166,792 | benign | N/A | SBP | 9.5E-11 | 84.7 |
| **379_2** | rs6271 | rs6271 | ENST00000393056.2 | *DBH*<br>p.Arg549Cys | 9 | 136,522,274 | possibly<br>damaging | tolerated | SBP | 1.2E-19 | 97.6 |
|  |  |  |  |  |  |  |  |  | DBP | 5.7E-35 | 100.0 |
| **394_1** | rs2236295 | rs2236295 | ENST00000373783.1 | *ADO*<br>p.Gly25Trp | 10 | 64,564,892 | possibly<br>damaging | tolerated | SBP | 2.8E-22 | 96.7 |
|  |  |  |  |  |  |  |  |  | DBP | 1.0E-28 | 89.4 |
| **402_3** | rs2274224 | rs2274224 | ENST00000371380.3 | *PLCE1*<br>p.Arg1575Pro | 10 | 96,039,597 | benign | tolerated | SBP | 9.0E-57 | 96.6 |
|  |  |  |  |  |  |  |  |  | DBP | 3.2E-44 | 93.4 |
| **406_5** | rs2484294 | rs1801253 | ENST00000369295.2 | *ADRB1*<br>p.Gly389Arg | 10 | 115,805,056 | benign | tolerated | DBP | 2.3E-74 | 89.3 |
| **412_3** | rs1133400 | rs1133400 | ENST00000368594.3 | *INPP5A*<br>p.Lys45Arg | 10 | 134,459,388 | benign | tolerated | SBP | 5.0E-15 | 77.8 |
|  |  |  |  |  |  |  |  |  | PP | 3.0E-10 | 86.1 |
| **414_7** | rs686722 | rs686722 | ENST00000418975.1 | *LSP1*<br>p.Arg12Trp | 11 | 1,891,722 | N/A | N/A | PP | 8.1E-19 | 82.6 |
| **417_10** | rs10770059<br>(SBP) | rs415895 | ENST00000318950.6 | *SWAP70*<br>p.Gln505Glu | 11 | 9,769,562 | benign | tolerated | SBP | 5.0E-47 | 96.5 |
|  | rs415895<br>(DBP) |  |  |  |  |  |  |  | DBP | 3.3E-69 | 96.8 |
| **432_4** | rs117874826 | rs117874826 | ENST00000540288.1 | *PLCB3*<br>p.Glu564Ala | 11 | 64,027,666 | benign | deleterious | SBP | 2.3E-11 | 100.0 |
| **434_1** | rs36027301 | rs36027301 | ENST00000265686.3 | *TCIRG1*<br>p.Arg56Trp | 11 | 67,809,268 | probably<br>damaging | deleterious | SBP | 6.0E-11 | 97.6 |
|  |  |  |  |  |  |  |  |  | DBP | 6.4E-12 | 77.8 |
| **447_7** | rs573455 | rs573455 | ENST00000278935.3 | *CEP164* | 11 | 117,267,884 | benign | tolerated | PP | 8.7E-34 | 100.0 |
|  |  |  |  | p.Gln1119Arg |  |  |  |  |  |  |  |
| **463_15** | rs1126930 | rs1126930 | ENST00000316299.5 | *PRKAG1* | 12 | 49,399,132 | benign | tolerated | PP | 4.6E-14 | 95.6 |
|  |  |  |  | p.Thr98Ser |  |  |  |  |  |  |  |
| **499_1** | rs17880989 | rs17880989 | ENST00000311852.6 | *MMP14* | 14 | 23,313,633 | benign | deleterious | DBP | 3.2E-12 | 100.0 |
|  |  |  |  | p.Met355Ile |  |  |  |  |  |  |  |
| **499_4** | rs365990 | rs365990 | ENST00000405093.3 | *MYH6* | 14 | 23,861,811 | benign | tolerated | SBP | 2.2E-12 | 90.8 |
|  |  |  |  | p.Val1101Ala |  |  |  |  |  |  |  |
| **523_5** | rs11854184 | rs11854184 | ENST00000559471.1 | *SECISBP2L* | 15 | 49,293,194 | benign | tolerated | DBP | 2.3E-10 | 86.8 |
|  |  |  |  | p.Val710Leu |  |  |  |  |  |  |  |
| **549_2** | rs55861648 | rs62622830 | ENST00000541260.1 | *C16orf93* | 16 | 30,770,950 | N/A | tolerated | PP | 1.2E-09 | 82.7 |
|  |  |  |  | p.Lys189Glu |  |  |  |  |  |  |  |
| **558_2** | rs62051555 | rs62051555 | ENST00000268489.5 | *ZFHX3* | 16 | 72,830,539 | probably<br>damaging | N/A | PP | 5.1E-25 | 89.9 |
|  |  |  |  | p.Gln2014His |  |  |  |  |  |  |  |
| **571_4** | rs1043809 | rs4924987 | ENST00000477478.2 | *B9D1* | 17 | 19,247,075 | benign | tolerated (LC) | DBP | 1.8E-16 | 77.2 |
|  |  |  |  | p.His143Tyr |  |  |  |  |  |  |  |
| **572_1** | rs704 | rs704 | ENST00000226218.4 | *VTN* | 17 | 26,694,861 | benign | tolerated | SBP | 1.9E-08 | 96.5 |
|  |  |  |  | p.Thr400Met |  |  |  |  |  |  |  |
| **573_1** | rs11080134<br>(SBP) | rs11080134 | ENST00000321990.4 | *ATAD5* | 17 | 29,161,503 | benign | deleterious | SBP | 1.8E-08 | 91.4 |
|  | rs9894838<br>(DBP) |  |  | p.Glu135Gly |  |  |  |  | DBP | 2.2E-08 | 79.5 |
| **576_26** | rs7406910<br>(SBP) | rs7406910 | ENST00000239165.7 | *HOXB7* | 17 | 46,688,256 | benign | tolerated | SBP | 1.1E-17 | 79.5 |
|  | rs9221 (PP) |  |  | p.Thr9Ala |  |  |  |  | PP | 5.1E-25 | 96.3 |
| **582_6** | rs34587622 | rs34587622 | ENST00000427177.1 | *SEPT9* | 17 | 75,398,498 | benign | tolerated (LC) | SBP | 6.2E-14 | 99.9 |
|  |  |  |  | p.Pro145Leu |  |  |  |  |  |  |  |
| **592_1** | rs61735998 | rs61735998 | ENST00000257209.4 | *FHOD3* | 18 | 34,289,285 | benign | deleterious<br>(LC) | PP | 2.0E-11 | 87.8 |
|  |  |  |  | p.Val647Phe |  |  |  |  |  |  |  |
| **606_7** | rs35918208 | rs2291516 | ENST00000393423.3 | *RGL3* | 19 | 11,508,177 | N/A | tolerated | DBP | 9.4E-11 | 85.7 |
|  |  |  |  | p.Arg621Cys |  |  |  |  |  |  |  |
| **606_9** | rs167479 | rs167479 | ENST00000393423.3 | *RGL3* | 19 | 11,526,765 | probably<br>damaging | deleterious | SBP | 8.7E-69 | 100.0 |
|  |  |  |  | p.Pro162His |  |  |  |  | DBP | 2.4E-79 | 100.0 |
|  |  |  |  |  |  |  |  |  | PP | 9.4E-21 | 100.0 |
| **608_1** | rs2108622 | rs2108622 | ENST00000221700.6 | *CYP4F2*<br>p.Val433Met | 19 | 15,990,431 | probably<br>damaging | deleterious | DBP | 1.1E-08 | 83.0 |
| **608_2** | rs4808045 | rs3745318 | ENST00000248071.5 | *KLF2*<br>p.Leu104Pro | 19 | 16,436,262 | benign | tolerated | DBP | 3.0E-14 | 89.1 |
| **610_2** | rs45522544 | rs45522544 | ENST00000357324.6 | *ATP13A1*<br>p.Glu556Lys | 19 | 19,765,499 | benign | tolerated | DBP | 3.1E-08 | 100.0 |
| **616_4** | rs34093919 | rs34093919 | ENST00000308370.7 | *LTBP4*<br>p.Asp752Asn | 19 | 41,117,300 | possibly<br>damaging | deleterious | PP | 2.8E-14 | 97.4 |
| **616_7** | rs1800470 | rs1800470 | ENST00000221930.5 | *TGFB1*<br>p.Pro10Leu | 19 | 41,858,921 | N/A | tolerated (LC) | PP | 1.9E-15 | 99.0 |
| **617_2** | rs7412 | rs7412 | ENST00000252486.4 | *APOE*<br>p.Arg176Cys | 19 | 45,412,079 | probably<br>damaging | deleterious | SBP | 2.0E-14 | 100.0 |
|  |  |  |  |  |  |  |  |  | PP | 2.8E-20 | 99.4 |
| **623_3** | rs35761929 | rs35761929 | ENST00000254958.5 | *JAG1*<br>p.Pro871Arg | 20 | 10,622,501 | benign | deleterious | DBP | 2.6E-18 | 99.1 |
| **636_1** | rs2229742 | rs2229742 | ENST00000400202.1 | *NRIP1*<br>p.Arg448Gly | 21 | 16,339,172 | probably<br>damaging | deleterious | SBP | 7.4E-16 | 100.0 |
|  |  |  |  |  |  |  |  |  | PP | 2.3E-11 | 99.7 |

*SNV, single nucleotide variant; Chr, chromosome; SIFT, Sorting Intolerant from Tolerant algorithm which predicts the effect of coding variants on protein function; Polyphen, Polymorphism Phenotyping tool predicts possible impact of an amino acid substitution on the structure and function of a human protein; Posterior probability, the SNV's accounted posterior probability of driving the blood pressure association under the annotation-informed prior.*

Fifteen missense variants were, in fact, identified to have a posterior probability of >99.9% of driving distinct BP association signals. These variants implicate several well characterised BP genes (including *SLC39A8* p.Ala391Thr; *ADRB2* p.Gly16Arg and p.Thr164Ile; and *DBH* p.Arg549Cys), but also highlight less well-established candidate genes, including *NRIP1, MMP14* and *PLCB3. NRIP1* is a regulator of the mineralocorticoid receptor, and *MMP14* is an endopeptidase with a key role in degrading components of the extracellular matrix and regulation of blood vessel stability^18^. EAWAS have identified missense variants in *PLCB3* to be associated with BP traits^7^, a gene that encodes an enzyme involved in intracellular signal transduction found to be increased in a mouse model of hypertension and hypertrophy^19^.

Several high-confidence missense variants implicate genes associated with kidney traits/disorders including *NCOA7* p. Ser399Ala, *LAMB2* p. Ala1765Thr and *NPHS2* p. Arg229Gln. *NCOA7* encodes the nuclear receptor coactivator 7, a vacuolar proton pumping ATPase (V-ATPase) interacting protein. It is highly expressed in the kidney with knock out mice observed to have lower BP^20^. *LAMB2* encodes beta chain isoform laminin, and mutations in this gene cause Pierson syndrome (OMIM# 609049), a congenital nephrotic syndrome, in which the phenotype includes hypertension^21^. Mutations in *NPHS2* cause steroid-resistant nephrotic syndrome^22^.

### Non-coding BP association signals map to trait-related transcription factor binding sites

Whilst the high-confidence missense variants identified have directly interpretable effects, the majority of the posterior probability of causality for BP trait associations maps to non-coding sequence. To explore these high-confidence non-coding SNVs, we firstly sought evidencefor enrichment of transcription factor binding site (TFBS) motifs. We interrogated sets of sequences obtained by expanding 10bp either side of these SNVs for each of the three traits (see **Methods**). This identified significant enrichment for 7 SBP, 10 DBP, and 5 PP TFBS motifs that were partially overlapping (Supplementary Table 8). The motif for *PAX2* was significant across all three traits (top corrected *P* = 2.8 × 10^−25^ for DBP), with this transcription factor involved in nephron development, as well as implicated in monogenic renal abnormalities^23^.

### Effector genes identified using gene expression in disease relevant tissues

To gain further insight into mechanisms through which non-coding association signals are mediated, where identification of the cognate effector gene is challenging^12^, we integrated genetic fine-mapping data with expression quantitative trait loci (*cis*-eQTL) in disease relevant tissues from the GTEx Consortium^24^. The tissues included were adipose, adrenal gland, artery, kidney cortex, heart, nerve, and brain. We observed convincing support for colocalization with eQTLs (**Methods)** for 96 SBP, 107 DBP and 84 PP signals (Supplementary Table 9). In total, 54 (56%), 58 (54%), and 41 (49%) of the signals colocalized with an eQTL for a single gene in at least one tissue. Across all traits, there was a total of 135 genes with tissue-specific colocalizations, of which 55 (41%) were in arterial tissues, 35 (26%) were in nerve or brain tissues, 21 (16%) were in adipose tissue, 13 (10%) were in heart, 9 (7%) were in adrenal, and 2 (1%) were in kidney.

Of the signals associated with all three BP traits, nine colocalized with an eQTL for a single effector gene. These were: *AGT* (brain cerebellum), *ARHGAP24* (tibial artery and aorta), *ARHGAP42* (tibial artery, aorta, and adipose), *CHD13* (aorta), *CTD-2336O2*.*1* (brain tissues), *FES* (tibial artery), *FGF5* (kidney cortex), *IGFBP3* (heart left ventricle), and *JPH2* (adrenal gland). Three genes (*AGT, ARHGAP42* and *IGFBP3*) have known or supporting data for a role in BP regulation. *AGT* encodes angiotensinogen, a substrate of the renin-angiotensinogen system – a key regulatory pathway^25^. *ARHGAP42* is selectively expressed in smooth muscle cells and modulates vascular resistance, and a knockout *Arhgap42* mouse model demonstrates salt mediated hypertension^26^. *IGFBP3*, which encodes the insulin growth factor binding protein 3, has data supporting association with BP and CVD phenotypes, and a knockout mouse model has increased ventricular wall thickness and shortened ST segment^27^. It also modulates insulin growth factor 1 (IGF-1)bioactivity with potential regulation of vascular tone *in-vivo* through NO release^28^. Additionally, there is a high-confidence missense variant implicating *IGFBP3*, highlighting distinct associations mediated by the same gene but through different underlying biological processes. Other colocalized effector genes demonstrate links to cardiovascular phenotypes (*FES*^29^, *FGF5*^30^, and *JPH2*^31^) but have not yet been functionally characterised.but demonstrate links to cardiovascular phenotypes.

We observed many individual loci with several distinct signals for each BP trait that colocalized with eQTLs for different genes. The genomic region on chromosome 12 encompassing *HDAC7, H1FNT, CCDC65, PRKAG1, AM186B, CERS5*, and *DIP2B*, spans 3.5Mb, and includes 11 signals for SBP, 12 for DBP and 5 for PP. Three signals colocalized with eQTLs and indicate two effector genes. One signal (associated with both SBP and DBP) colocalized with an eQTL for *CACNB3* (adipose, tibial nerve and artery), which encodes a regulatory beta subunit of the voltage-dependent calcium channel. The regulatory subunit of the voltage-gated calcium channel gives rise to L-type calcium currents^32^. A *CACNB3* knock-out mouse model has a cardiovascular phenotype that includes abnormal vascular smooth muscle cell hypertrophy, increased heart weight and increased SBP and DBP^33^. A second signal, associated with DBP colocalized with an eQTL for *RP4-605O3*.*4* (heart left ventricle), and a third signal (associated with SBP) colocalized with an eQTL in brain and heart left ventricle for this predicted gene.

A genomic region on chromosome 17 spanning 6.4Mb, which encompasses associations reported in several previous BP GWAS^3,7,34,35^,includes 19 SBP, 16 DBP and 15 PP signals (Supplementary Table 2). Colocalization of signals with eQTLs implicates six effector genes (*DCAKD, NMT1, RP11-6N17*.*4, PNPO, PRR15LA* and *ZNF652*). Three independent signals colocalized with eQTLs for *NMT1* in brain. The *NMT1* gene encodes N-myristoyltransferase, which catalyses the transfer of myristate from CoA to proteins, and there is no clear association with cardiovascular disease. However, the Malacards database indicates an association with Patent Foramen Ovale, a common post-natal defect of cardiac atrial septation^36^. One DBP signal colocalized with an eQTL for *DCAKD* in adipose and nerve tissues. PP signals colocalized with eQTLs for *RP11-6N17*.*4* and *PNPO* in brain tissues. *PNPO* encodes pyridoxamine 5’-phosphate oxidase, an enzyme in the rate limiting step in vitamin B6 synthesis. Deficiency of PNPO primarily results in seizures, with many systemic symptoms including cardiac abnormalities^37^. We also observed a SBP signal that colocalized with an eQTL for *PRR15LA* in tibial artery, and a signal associated with both with SBP and DBP that colocalized with an eQTL for *ZNF652* in adipose tissue.

At a second genomic region on chromosome 17 encompassing *MRC2, ACE, PECAM1*, and *MILR1*, we observed four signals for SBP, three for DBP and four for PP (Supplementary Table 2), of which three signals colocalized with different genes across multiple tissues (Figure 3). One SBP signal colocalizes with an eQTL for *MRC2* in tibial artery. *MRC2* encodes the mannose receptor C type 2 and plays a role in extracellular matrix remodelling^38^. A signal associated with both SBP and DBP colocalized with an eQTL for *ACE* in kidney, adipose, and brain tissues. *ACE* encodes the angiotensin-converting enzyme, a central component of the renin–angiotensin– aldosterone system^39^. A third SBP signal colocalized with an eQTL for two genes across several tissues: *DDX5* (arteries, brain and tibial nerves) and *CEP95* (tibial nerve and arteries). These genes have little prior association with cardiovascular phenotypes. *DDX5* encodes DEAD-Box Helicase 5, which is thought to be a coregulator of transcription or splicing, and recent data indicates a role in smooth muscle cell protection and neointimal hyperplasia^40^. Homozygous *Ddx5* knockout mice die at embryonic day 11.5 and demonstrate blood vessel abnormalities. There is little information on *CEP95*, which encodes centrosomal protein 95, although differential gene expression was observed in spontaneously hypertensive rats^41^.

**Figure 3.**
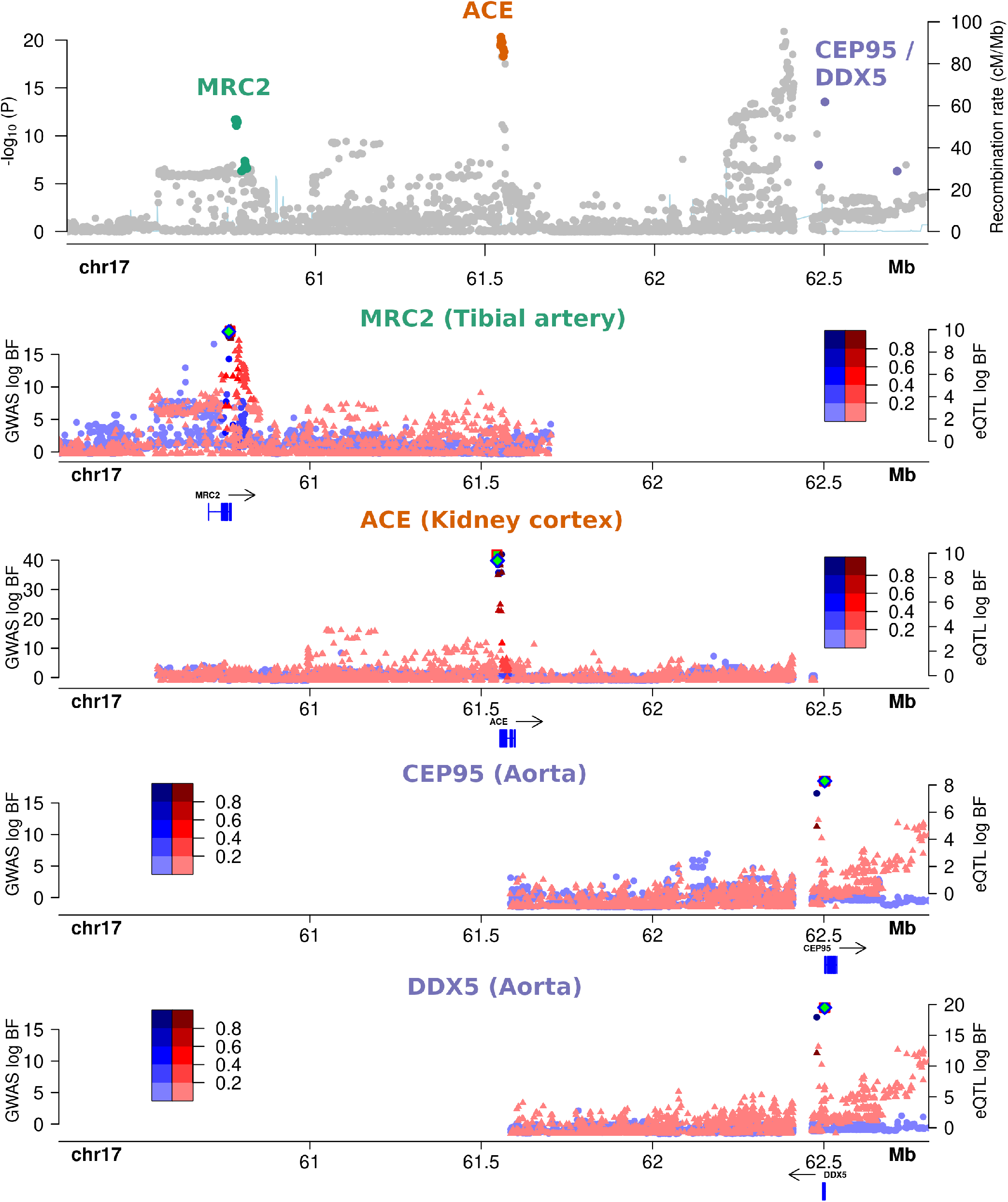
Colocalization between GWAS signals for systolic blood pressure and multi-tissue expression data at locus ID 580 on chromosome 17. The top panel shows the unconditional GWAS data of the genomic region at chromosome 17 (60.2Mb – 62.9Mb) for systolic blood pressure. The lower four panels show the log annotation informed Bayesian factors of the conditional GWAS signal (blue, left axis) and gene expression data from GTeX eQTL data (red, right axis). Three distinct annotation informed signals colocalized with gene expression data from: *MRC2* (second panel) in tibial artery tissue, *ACE* (third panel) in kidney cortex tissue, and *CEP95* and *DDX5* in aortic artery tissue (bottom panels). The x-axis shows the physical position on the chromosome (Mb) and the y-axes show the log annotation informed Bayesian factor from the GWAS (left axis) and the gene expression data (right axis). The intensity of the color indicates the linkage disequilibrium with respect to the sentinel GWAS SNP (blue) or top eQTL SNP (red).

### Identification of effector genes using promotor-centered long-range chromatin interactions in disease relevant tissues

To explore possible long-range enhancer influence on specific target genes, we integrated genetic fine-mapping data with potential functional CREs identified to target the promoters of well-annotated protein-coding genes via long-range chromatin interactions (capture Hi-C data from Jung et al.^42^). Promoter interactions and candidate genes were identified for 629 signals at 366 genomic regions (RegulomeDB Score ≤ 3) across adrenal gland, dorsolateral prefrontal cortex, hippocampus, aorta, left ventricle, right ventricle, and fat (Supplementary Table 10). We observed signals at 13 genomic regions that included 99% credible set variants with regulatory potential across SBP and DBP, for which several potential target genes of the regulatory variants were indicated. At five signals, one gene was indicated in a single tissue: *ACTRT2* (dorsolateral prefrontal cortex), *ARMC4* (right ventricle), *AP001024*.*1* (hippocampus), *TBX3* (aorta) and *YES1* (hippocampus). At other genomic regions, many genes in one tissue were indicated: *HOXA5, HOXA6* and *HOXA3* (adrenal gland), *RP11691N7*.*6, C11orf31, TIMM10, CLP1, YPEL4, ZDHHC5, FAM111A, TIMM10, AP001350*.*1, GLYATL2, GLYATL1* (hippocampus, dorsolateral prefrontal cortex), and *ABHD17C, MESDC2* (two brain tissues). At three signals, more than one gene in more than one tissue were highlighted, such as *GPR124, DDHD2, FGFR1, PPAPDC1B, LETM2, TACC1* (two brain tissues, adrenal gland, aorta, and right ventricle). Candidate genes at three signals have existing functional data supporting an association with BP or cardiovascular traits: *HOXA3, ADM* and *TBX3*^*27,43,44*^.

We next explored whether signals that colocalized with eQTLs for effector genes overlapped with those implicated by Hi-C predicted promoter interactions. We focused on the 80 signals that have support for colocalization with eQTLs in relevant tissues across all traits (Supplementary Table 11). For 34 signals, the effector gene indicated by Hi-C was the same as that identified via colocalization with the eQTLs, and for 15 of these signals the same gene and tissue was implicated (Supplementary Table 12). The 15 candidate genes were: *AKR1B1, ASAP2, COL27A1, IRF5, MAP1B, MRPS6, MXD3, RAD52, RERE, RNF130, SLC5A3, SLC20A2, TNS3, TRIOBP*, and *USP36*. A review of the 15 genes indicates knock-out mouse models of three effector genes (*COL27A1, RERE*, and *SLC20A2*) have cardiovascular abnormalities – these genes have not previously been highlighted as potential candidate genes for hypertension (Supplementary Table 12).

To explore our Hi-C predicted promoter interactions more broadly, we additionally probed our results to see whether there was also support for these potential CREs to target the same effector gene through a completely different prediction methodology from the recent EpiMap analysis^45^. This method is based on an active chromatin correlation with target gene expression. Several physiologically relevant candidate effector transcripts were highlighted, where the two methods predicted the same target for the same SNV in the same tissue or organ (See Supplementary Table 13). Potential targets included genes previously identified as highly plausible trait-related candidates from previous analyses, including *CLIC4, TNS1*, and *FERMT2*^46^. Target genes with presently unknown potential roles in SBP and DBP pathophysiology were also identified. These included two active CREs found in brain-related tissue, which would be of interest to explore for additional activity in potentially more physiologically relevant non-assayed tissues: *SHMT1*, the serine hydroxymethyltransferase 1 enzyme involved in folic acid metabolism associated, although inconsistently, with hypertension-related stroke^47^ and *PLXNB2*, the Plexin-B2 transmembrane receptor that has an identified role in the developing kidney^48^. For PP this comprised, amongst others, some interesting target genes including *MYH11*, the smooth muscle myosin heavy chain 11 gene, with CRE activity in aortic tissue. Mutations within this gene lead to an autosomal dominant aortic aneurysm and dissection disorder (*AAT4*, OMIM #132900) with altered aortic stiffness^49,50^. Also, *COL6A3*, which encodes the alpha-3 chain of type VI collagen, was a target identified in heart tissue. This collagen gene is important in the developing mammalian heart^51^ and is a causative gene in monogenic myopathy and dystonia diseases (OMIM: #158810; #254090; #616411)^52^.

### Consolidated effector gene evidence

Using complementary fine mapping and computational approaches (high-confidence missense, colocalised eQTLs and Hi-C interactions) we identified 956 candidate genes for SBP, 900 candidate genes for DBP and 773 candidate genes for PP with at least one line of evidence indicating a putative effector gene (Supplementary Tables 14-16).

We next looked for additional supportive evidence for each gene by combining information from mouse model data, human cardiovascular and renal phenotypes, and differential gene and protein expression across cardiovascular tissues (Methods). We selected as consolidated effector genes those which had two or more additional lines of evidence. In total, 197 SBP, 184 DBP and 180 PP genes were identified (Supplementary Tables 17-19), which together reflect 394 unique candidate genes.To gain insights into the biological role of these consolidated evidence effector genes for each BP trait we performed gene-set enrichment analyses. We found significant enrichment for 390, 333, and 299 gene ontology (GO) biological processes for SBP, DBP, and PP respectively (following removal of redundant processes, see Methods). There were 629 unique GO ID terms across the three traits, with 153, 106 and 103 unique to SBP, DBP and PP respectively. In total, 141 pathways were associated with two BP traits and 126 pathways with all three BP traits (Supplementary Tables 20-22). Some of the pathways associated with all three BP traits included: blood vessel remodelling, regulation of the immune system, circulatory and renal system processes, sodium ion transport and ion homeostasis, aging, smooth muscle cell migration, lipid metabolism processes and cytoskeletal organisation – all processes previously highlighted as important in BP control. The most significant SBP unique processes included: neurogenesis (*P* =1.9 × 10^−17^), heart development (*P* = 1.2 × 10^−11^) and regulation of cell death (*P* = 3.4 × 10^−7^); for DBP, embryonic organ development (*P* = 5.0 × 10^−11^) and positive regulation of RNA biosynthetic processes (*P* = 1.1 × 10^−9^); and, for PP, muscle organ development (*P* = 5.1 × 10^−9^) and trabecular formation (*P* = 0.007) and morphogenesis (*P* = 0.004).

### Drug target identification and repositioning opportunities

We assessed the druggability of the consolidated candidate effector genes for each BP trait via the druggable genome dataset from Finan et al.^53^ (Supplementary Table 23, see Methods). We observed DBP to have a greater number of candidate effector genes that encode proteins that are the main drug targets for anti-hypertensive medications (*ACE, ADRA1A, ADRB1* and *NR3C2*), compared with SBP (*ACE*) and PP (none). For several effector genes that are targets of existing drugs, there was support as potential therapeutic targets for hypertension (e.g., *AKR1B1, PDE3A* and *MAP2K1*). *AKR1B1* (aldo-keto reductase family 1 member B) is a target of aldose reductase inhibitors that have been investigated for use in diabetes and also have effects on BP^54^. *PDE3A* (phosphodiesterase 3A) is a target for hypertension with bracydactyly, a rare autosomal dominant disorder and there is recent data indicating several common variant associations also in the general population^7,55^. *PDE3A* is targeted by several existing drugs, including Cilostazol (peripheral vascular disease), Levosimendan for intravenous therapy for acutely decompensated heart failure and Enoximone (pulmonary hypertension). There are no data currently indicating the use of *PDE3A* inhibitors for hypertension, however a recent study suggests activation of *PDE3A* in the heart may protect it from hypertrophy and failure^56^. *MAP2K1* (*MEK1*) is a target of anti-neoplastic agents (*MAP2K1* is altered in 1% of lung and head and neck squamous cell carcinomas)^57,58^.The *MAPK* pathway is well recognised in BP control and p38-MAPK inhibition has been considered previously as a therapeutic target (which *MAP2K* activates). Our study applied rigorous multiple evidence methodology employing differing datasets from the Evangelou et a to derive a consolidated effector gene list.^3^ This impacted the identification of potential therapeutic targets as we were not able to replicate in our analysis the five candidate genes (*CA7, CACNA1C, CACNB4, PKD2L1* and *SLC12A2*) reported in their paper as the target of anti-hypertensive drug classes. However, *PKD2L1* did show individual Hi-C evidence as bring the effector gene at the locus.

To further ascertain drug repositing opportunities, we tested for enrichment of these consolidated candidate effector genes in clinical indication categories. We observed significant enrichment of gene sets for cardiovascular and renal conditions (Supplementary Table 24), with the results supporting the findings from interrogation of the Finan et al. druggable genome database.

## Discussion

Strongly replicated human genetic associations with BP traits have been identified over the last decade. In this study, we have used a robust contemporary fine mapping pipeline to advance from these initial broadly associated genomic regions to the identification of hundreds of plausible previously-unreported effector genes (Figure 4). These candidates are now excellent targets for future focused functional validation.

**Figure 4.**
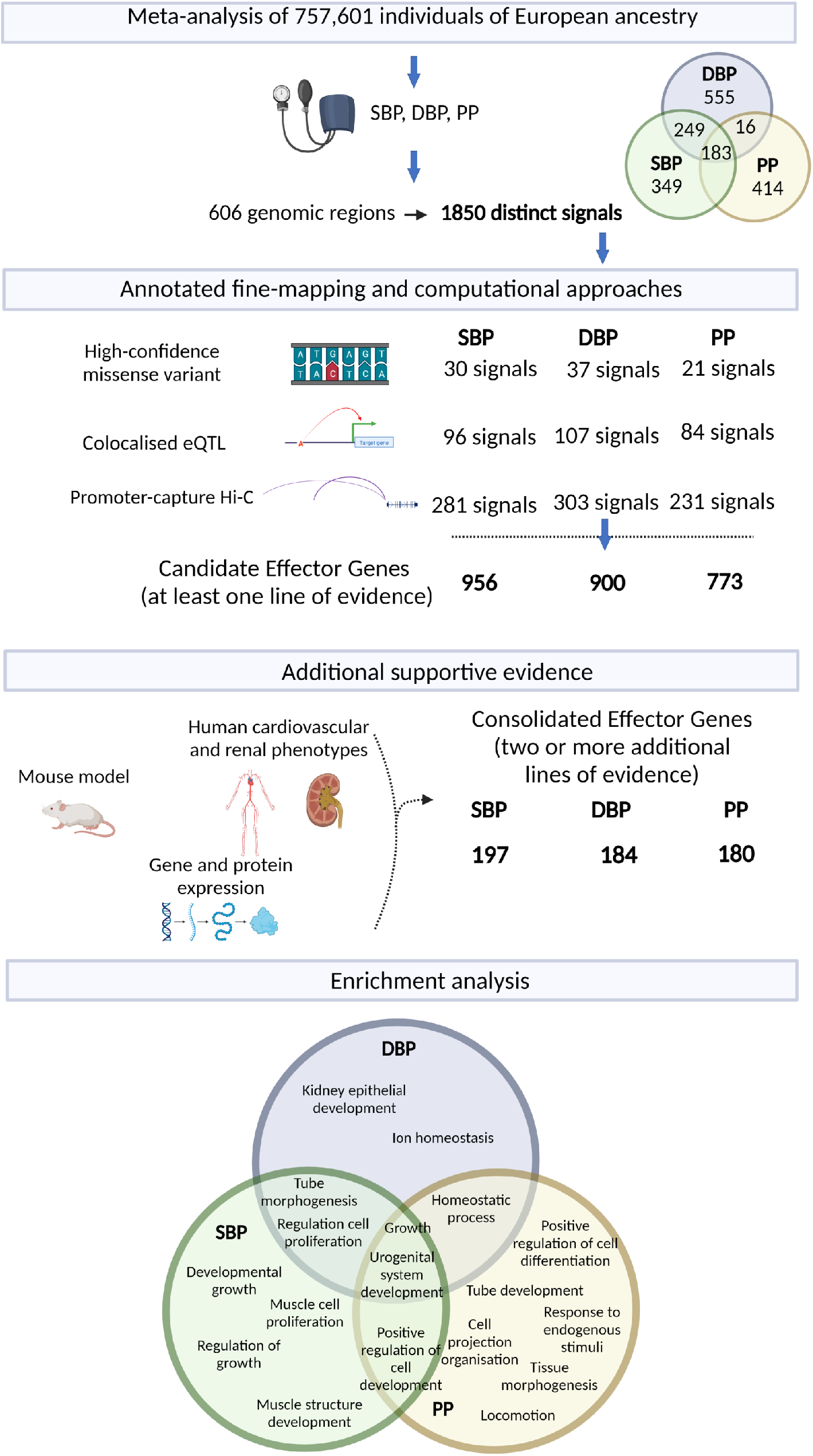
BP fine mapping results and consolidated effector genes. Overview of results from the fine-mapping pipeline for SBP, DBP and PP. For each trait the number of signals are shown following annotation as missense, eQTL or regulatory, and the number of candidate effector genes based on at least one line of evidence. Using additional evidence from mouse models, human cardiovascular and renal phenotypes and gene/protein expression the list of effector genes was reduced – these are the consolidated effector genes. The consolidated effector genes were used as input to FUMA (methods) for pathway enrichment analysis. Results are shown for GO terms after removal of similar terms using the Reduce and Visualise Gene Ontology (REVIGO) web application (dispensibility cut off <0.3), and only pathways which had adjusted *P* values <10^−10^ were included. Created with Biorender.com

We were able to localise approximately a quarter of all associations across all three BP traits to a single causal variant with >75% posterior probability. Of these high-confidence SNVs, 65 were missense variants, including 20 for two BP traits, and one in *RGL3* for all three traits. For the high confidence non-coding and potentially *cis*-regulatory variants, we employed pathogenic tissue-specific expression and chromatin conformation to identify their target genes. Of these SNVs, ∼100 per trait colocalized with *cis*-eQTLs. Plausible effector genes included the well-known Angiotensin (*AGT*) and angiotensin converting enzyme (*ACE*), also more recently described genes from GWAS with functional data including sodium/potassium-transporting ATPase subunit beta-1 (*ATP1B1*) and Rho GTPase activating protein 42 (*ARHGAP42*). Other possible but less well functionally evaluated genes identified through eQTL analysis included *CDH13, FES, FGF5*, and *JPH2*.

We identified many loci with multiple complex signals within the same genomic region affecting different genes in different tissue types. Also, of note that whilst we observed high-confidence missense variants in kidney genes, as well as an enrichment for non-coding variants to overlap a nephron developmental TFBS, we identified only a very small proportion of eQTL colocalization in this tissue (*FGF5* and *ACE* and the lncRNA *AC021218*.*2*). This may reflect reduced power due to the relatively smaller numbers in GTEx for kidney than other tissues^24^. Using over 400 human kidneys and the same ICBP+UKBB GWAS dataset, Eales and colleagues reported nearly 31% of BP associated variants contained kidney eSNPs^59^. These results strongly emphasise the importance of access to larger tissue banks for robust identification of all possible effector genes.

Focusing on the overlap between the eQTL and Hi-C results in disease relevant tissues identified a subset of 15 target genes identified in the same tissue. Of these, the genes *COL27A1, RERE*, and *SLC20A2* also had supportive mouse data. Furthermore, there is evidence that target genes consistently predicted across multiple methods are the most robust^60^. We also explored overlap with the recent EpiMap^61^, highlighting, amongst others, *MYH11* and *COL6A3* as strong effector candidates for PP. In total, our pipeline identified consolidated evidence effector genes for ∼20% of BP association signals (196 genes for 865 SBP associations; 184 genes for 904 DBP associations; and 184 genes for 697 PP associations), an overview of the main findings from the study are illustrated in Figure 4. Of the plausible BP genes, 14% were identified to be drug targets, and several of these have good support for potential repurposing for BP control.

The main strength of this work is that it combines robust GWAS associations, derived from a powerfully large dataset, with comprehensive genetic annotation and tissue-specific epigenomic maps derived from the Epigenomics Roadmap consortium. In the exploration of the putative functional non-coding variants, a further strength was that we were able to benefit from the expanded GTEx dataset^24^, publicly available promoter capture Hi-C data in pathogenically relevant tissues^42^, as well as exploration of target gene prediction overlap with EpiMap^45^. These analyses identified many biologically plausible effectors genes.

Current weaknesses are the lack of population diversity in our GWAS dataset, as these are comprised of associations only from European ancestry individuals. Consequently, they will be missing population-specific findings, as have been identified in other common diseases^62^. Furthermore, this lack of diversity is not only limited to the genetic findings. The epigenomic maps, whilst being derived from a breadth of cell-types giving good representation of strong tissue-specific regulatory differences, are within each cell-type drawn from very small numbers. Therefore, they lack detail regarding potential population variation in these functional units^63^. Another weakness is that whilst benefitting from dense genotyping and imputation of common SNVs, this is not exhaustive in capturing all the potential phenotypically associated genetic variation within each locus. This will miss the possible impact of rare SNVs, as well as any poorly tagged larger variants (copy number variants, short tandem repeats, inversions, *etc*.). Furthermore, these large variants may themselves facilitate functional epigenomic variation^64^. Future exploration of the phased interplay of genetic and epigenetic allelic elements by advancing long-read technologies will help to fill in these gaps^65^.

In conclusion, we have identified plausible causal common genetic variants enriched in BP pathways. Their investigation through experimental biosystems will not only improve functional understanding of the biology of BP and its pathogenesis, but also potentially enable novel preventative and therapeutic opportunities.

## Methods

### Study data and detection of distinct association signals

We utilised previously reported GWAS meta-analyses of blood pressure traits in up to 757,601 individuals of European ancestry from the ICBP and UK Biobank^3^ (ICBP+UKBB). Each contributing GWAS had been imputed up to reference panels from the 1000 Genomes Project ^9,10^ and/or Haplotype Reference Consortium^11^. After quality control, meta-analysis association summary statistics for SBP, DBP and PP were reported for 7,088,121, 7,160,657 and 7,088,842 SNVs, respectively. We began by considering autosomal lead SNVs that have been reported at genome-wide significance (variable threshold according to study design) for SBP, DBP or PP in previously published GWAS of blood pressure traits, which we have collated and are summarised in the recent review by Magavern and colleagues^8^. We initially defined genomic regions as mapping 500kb up- and down-stream of each lead SNV. However, where genomic regions overlapped, they were combined as a single genomic region to account for potential LD between previously reported lead SNVs. Genomic regions that did not attain genome-wide significance (*P* < 5×10^−8^) in the ICBP+UKBB meta-analysis for any BP trait were not considered for downstream interrogation. We then performed approximate conditional analyses using GCTA-COJO^66^ to detect distinct association signals at each genomic region for each BP trait separately, using European ancestry haplotypes from the 1000 Genomes Project (Phase 3, October 2014 release)^9^ as a reference for LD. Within each genomic region variants attaining genome-wide significance (*P* < 5×10^−8^) in the joint GCTA-COJO model were selected as index SNVs for distinct association signals.

We next assessed the evidence that distinct association signals for SBP, DBP and PP were shared across multiple BP traits. At each genomic region distinct association signals for two traits were considered to be the same if: (i) the index SNVs were the same for both traits; (ii) the index SNVs were colinear in the joint GCTA-COJO models for each trait after including the index SNV for the other trait in the model; or (iii) the P-value of the index SNV for one trait increased to P > 0.05 after including the index SNV for the other trait in the model, and the P-value of the index SNP for the other trait increased to P > 0.0001 for the corresponding reciprocal conditioning.

### Enrichment of BP associations for genomic annotations

We used fGWAS^67^ to identify genomic annotations enriched for SBP, DBP or PP association signals. We considered a total of 253 functional and regulatory annotations derived from: (i) genic regions (protein coding exons, 3’ UTRs and 5’ UTRs) as defined by the GENCODE Project^13^; and (ii) chromatin state predictions of promoters and enhancers across 125 tissues from the Roadmap Epigenome Consortium^15^ via Epilogos (http://compbio.mit.edu/epilogos/). For each BP trait separately, we used a forward-selection approach to derive a joint model of enriched annotations. At each iteration, we added the annotation to the joint fGWAS model that maximised the improvement in the penalised likelihood. We continued until no additional annotations improved the fit of the joint model (P<0.00020, Bonferroni correction for 253 annotations).

### Fine-mapping distinct association signals for BP traits

For each trait, we began by approximating the Bayes’ factor (BF), *A*_*ij*_, in favour of association of the *j*th SNV at the *i*th distinct association signal using summary statistics from the ICBP+UKBB meta-analyses. Specifically,

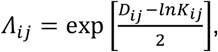

where 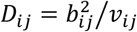, and *b*_*ij*_ and *v*_*ij*_ are the allelic log-OR and corresponding variance, respectively, across *K*_*ij*_ contributing GWAS to the ICBP+UKBB meta-analysis (here *K*_*ij*_ = 2)^68^. At genomic regions with a single association signal, *b*_*ij*_ and *v*_*ij*_ were taken from the unconditional meta-analysis. However, for genomic regions with multiple association signals, *b*_*ij*_ and *v*_*ij*_ were taken from the joint GCTA-COJO model, conditioning on the index SNVs for all other signals at the locus. The posterior probability for the *j*th SNV at the *i*th distinct signal, was then given by *n*_*ij*_ ∝ *y*_*j*_*A*_*ij*_, where *y*_*j*_ is the relative prior probability of causality for the *j*th SNV. We considered an annotation-informed prior model, for which

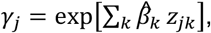

where the summation is over the enriched annotations, 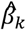-is the estimated log-fold enrichment of the *k*th annotation from the final joint fGWAS model, and *z*_*jk*_ is an indicator variable taking the value 1 if the *j*th SNV maps to the *k*th annotation, and 0 otherwise. Finally, we derived a 99% credible set^69^ for the *i*th distinct association signal by: (i) ranking all SNVs according to their posterior probability *n*_*ij*_; and (ii) including ranked SNVs until their cumulative posterior probability attains or exceeds 0.99.

### High Confidence SNV Gene Set Enrichment Analysis

Genomic Regions Enrichment of Annotations Tool (GREAT) v4.0.4^16^ was used to explore the high confidence SNVs potential biological impact. The default GREAT association parameters for gene regulatory domains (Proximal 5 kb upstream, 1 kb downstream, plus Distal up to 1 Mb) were used and curated regulatory domains included. Input was SNV BED files for each of the three traits (SBP n = 208, DBP n = 224 and PP n = 158). GREAT analysis included gene ontology (GO) Biological Processes, Human Phenotype, Mouse Phenotype and Knockout data.

### Functional annotation

We use Variant-effect predictor (VEP) analysis to identify missense variants and queried their overlap with high confidence causal variants from the credible set analysis (https://grch37.ensembl.org/Homo_sapiens/Tools/VEP)^70^.

### Transcription Factor Binding Motif Analysis

We used the Transcription Factor Affinity Prediction (TRAP) v3.0.5^71^ multiple sequences option to explore any enrichment for Transcription Factor (TF) binding motifs within the high confidence non-coding variants for each of the three traits (SBP n= 178; DBP n=187; and PP n =137). Sequences around each non-coding SNV were expanded to +/-10bp (via AWK) and the FASTA sequence extracted (hg19) via the BEDtools v2.30.0 command getfasta^72^. The Transfac 2010.1 Vertebrate matrix set was interrogated with human_promoter set as background model and the results were required to pass a Benjamini-Hochberg multiple-testing correction.

### Colocalization with gene expression data

We performed a Bayesian statistical procedure to assess whether our annotation informed GWAS results were colocalised with eQTL results. We used all available eQTL tissues relevant for blood pressure (adipose, adrenal gland, artery, kidney cortex, heart, nerve, and brain) from the publicly available eQTL results from GTEx version 8^24^. The annotation informed BF in favour of association of the *j*th SNV at the *i*th distinct association signal was defined as:

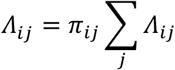

In this expression, *n*_*ij*_ is the annotation-informed posterior probability, and *A*_*ij*_ is the BF defined above. GWAS results were lifted from hg19 to hg38 used lift Over software^73^ [to allow direct comparison with the hg38 eQTL data. We undertook colocalization using the annotation informed BF using the COLOC software package in R^74^, only for those signals for which a 99% credible set variant was the lead eQTL SNV.

### Long-range chromatin interaction (Hi–C) analyses

We identified potential target genes of regulatory SNVs using long-range chromatin interaction (Hi–C) data from tissues and cell types relevant for blood pressure regulation (adrenal gland, left and right ventricles, hippocampus, and cortex)^42^. Hi–C data is corrected for genomic biases and distance using the Hi–C Pro and Fit-Hi-C pipelines according to Schmitt et al. (40 kb resolution—correction applied to interactions with 50 kb-5Mb span)^75^. We selected the most significant promoter interactions for all potential regulatory SNPs (RegulomeDB score ≤3) that were included in the 99% credible sets and report the interactors with the SNPs of highest regulatory potential to annotate the loci.

### Collation of evidence for effector BP genes

A full list of candidate effector genes for each BP trait was collated from the results of our fine-mapping pipeline and computational approaches. A gene was indicated for a signal if there was support from a coding and high confidence variant in the gene at the locus, or if the gene was indicated from eQTL colocalization or Hi-C analyses. To refine the list of putative candidate effector genes we next collated additional information for each gene using data from GeneCards (https://genealacart.genecards.org). This included the following: 1) a mouse model from Mouse Genome Informatics (MGI) which has a cardiovascular or renal phenotype. 2) A cardiovascular, vascular or renal phenotype described for the candidate gene in the Human Phenotype Ontology database 3) Differential RNA expression of the candidate gene in the GTEx database in cardiovascular, vascular or renal tissues, only genes with fold changes >4 in a tissue were selected. 4) Differential protein expression of the candidate gene based on 69 integrated normal proteomics datasets in HIPED (the Human Integrated Protein Expression Database). Genes with a fold change value of >6 and protein abundance value of >0.1 PPM in an anatomical were selected. The top effector candidate genes for each BP trait were selected if there were at least 2 additional lines of evidence.

### Effector gene pathway analysis

We used the Gene2Function analysis tool in FUMA (v1.4.0) to perform geneset enrichment and identify significantly associated GO terms and pathways^76^. Hypergeometric tests were performed to test if genes were over-represented in any predefined gene set and multiple testing correction was performed per category. The gene sets used are from MsigDB, WikiPathways and genes from the GWAS-catalog. The analysis included the putative top effector genes only. The analysis was done for all BP traits and we report results with adjusted p-values of <0.05. Redundant GO terms were removed using the Reduce and Visualize Gene Ontology (REVIGO) web application^77^. REVIGO uses a hierarchical clustering method to remove highly similar terms, incorporating enrichment *P*-values in the selection process. Default settings (dispensability cut off <0.7) were used in this analysis.

### Druggability of prioritised effector genes

To identify candidate druggable targets, a look-up was performed in a previously published database of the druggable genome developed by Finan et al.^53^ This list contains protein-coding genes catagorised into three tiers: Tier 1 are targets of approved drugs and some drugs in clinical development, including targets of small molecules and biotherapeutics; Tier 2 are proteins closely related to drug targets or associated with drug-like compounds (≥50% shared protein sequence identity); Tier 3 includes extracellular proteins and members of key drug target families in Tier 1 (e.g., G protein-coupled receptors). To identify potential opportunities for drug repurposing, a look-up of each BP candidate gene was performed in Tier 1 to identify existing drug targets (https://www.genome.jp/kegg/genes.html). Primary targets of antihypertensives were also identified using the KEGG drug database (https://www.genome.jp/kegg/drug/). The open targets database was subsequently interrogated to identify disease associations with each gene, to identify potential overlap that could indicate promising drug targets. Target, drug and disease association data was downloaded from the platform (https://platform.opentargets.org/downloads). Open targets calculates association scores to capture the data type (e.g., gene level) and source, to aggregate evidence for an association, by calculating the harmonic sum using a weighted vector of data source scores. This sum is divided by the maximum theoretical value, resulting in a score between 0 and 1. To identify enrichment of candidate effector genes in clinical indication categories and potentially re-positional drugs, we utilised the Genome for REPositioning drugs (GREP) software^78^. GREP performs a series of Fisher’s exact tests, to identify enrichment of a gene-set in genes targeted by a drug in a clinical indication category (Anatomical Therapeutic Chemical Classification System [ATC] or International Classification of Diseases 10 [ICD10] diagnostic codes).

## Acknowledgements

This research was supported by the NIHR Barts Cardiovascular Research Centre and the NIHR Manchester Biomedical Research Centre (NIHR203308). J.R. acknowledges funding from the European Union’s Horizon 2020 Research and Innovation Programme under the Marie Sklodowska-Curie grant agreement number 786833, from the European Union-NextGenerationEU, and from Grant PID2021-128972OA-I00 funded by MCIN/AEI/ 10.13039/501100011033. W.J.Y. recognises the National Institute for Health Research (NIHR) Integrated Academic Training programme, which supports his Academic Clinical Lectureship post.

## Ethics declarations

The authors declare no competing interests.

## Author Contributions

S.vD., J.R., W.J.Y., C.G.B., A.P.M., P.B.M designed the study. S.vD., J.R., W.J.Y., K.J.O., F.A., M.J.A.Y.A., C.G.B.., A.P.M., P.B.M. performed analyses. S.vD., J.R., W.J.Y., C.G.B., A.P.M., P.B.M. drafted the manuscript, and all authors provided critical revisions.

