## Supplemental information for "Integration of genetic fine-mapping and multi-omics data reveals candidate effector genes for hypertension"

**Van Duijvenboden et al**

### (A) Supplementary Figure Legends

#### Supplementary Figure 1

Flow chart illustrating the steps followed in the fine-mapping analysis of blood pressure traits.

* eQTLs in artery, adipose, heart, brain and kidney tissues (GTEx, v8); † adrenal gland, left and right ventricles, brain and fat tissues.

SBP, systolic blood pressure; DBP, diastolic blood pressure; PP, pulse pressure; GWAS, genome-wide association study; fGWAS, functional genome-wide association study; VEP, variant effect predictor; eQTL, expression quantitative locus.

#### Supplementary Figure 2

Results from genomic enrichment annotation for each BP trait. Estimate and 95% confidence interval of the enrichment at the most significant annotations for SBP, DBP and PP, calculated using functional GWAS.

UTR, untranslated region; HUVEC, human umbilical vein endothelial cells; NHDF-Ad, neonatal human dermal fibroblasts adult; TSS, transcription start site; SBP, systolic blood pressure; DBP, diastolic blood pressure; PP, pulse pressure.

#### Supplementary Figure 3

A: Systolic Blood Pressure – (GREAT analysis)

The high confidence variants were tested for enrichment via the Genomic Regions Enrichment of Annotations Tool (GREAT v4.0.4, PMID: 20436461) for GO Biological Processes, Human Phenotype, Mouse Phenotype and Knockout datasets – see Methods. Figures display –log(10) Binomial p values.

A: Diastolic Blood Pressure – (GREAT analysis)

The high confidence variants were tested for enrichment via the Genomic Regions Enrichment of Annotations Tool (GREAT v4.0.4, PMID: 20436461) for GO Biological

Processes, Human Phenotype, Mouse Phenotype and Knockout datasets – see Methods. Figures display –log(10) Binomial p values.

B: Pulse Pressure – (GREAT analysis)

The high confidence variants were tested for enrichment via the Genomic Regions Enrichment of Annotations Tool (GREAT v4.0.4, PMID: 20436461) for GO Biological Processes, Human Phenotype, Mouse Phenotype and Knockout datasets – see Methods. Figures display –log(10) Binomial p values.

### (B) International Consortium of Blood Pressure (ICBP) - Authors & Contributors

From the 2018 Nature Genetics publication: "Genetic analysis of over one million people identifies 535 new loci associated with blood pressure traits."

Evangelos Evangelou[1,2], Helen R Warren[3,4], He Gao[1,5], Georgios Ntritsos[2], Niki Dimou[2], Tonu Esko[16,17], Reedik Mägi[16], Lili Milani[16], Peter Almgren[18], Thibaud Boutin[19], Stéphanie Debette[20,21], Jun Ding[22], Franco Giulianini[23], Elizabeth G Holliday[24], Anne U Jackson[25], Ruifang Li-Gao[26], Wei-Yu Lin[27], Jian'an Luan[28], Massimo Mangino[29,30], Christopher Oldmeadow[24], Bram Peter Prins[31], Yong Qian[22], Muralidharan Sargurupremraj[21], Nabi Shah[32,33], Praveen Surendran[27], Sébastien Thériault[34,35], Niek Verweij[17,36,37], Sara M Willems[28], Jing-Hua Zhao[28], Philippe Amouyel[38], John Connell[39], Renée de Mutsert[26], Alex SF Doney[32], Martin Farrall[40,41], Cristina Menni[29], Andrew D Morris[42], Raymond Noordam[43], Guillaume Paré[34], Neil R Poulter[44], Denis C Shields[45], Alice Stanton[46], Simon Thom[47], Gonçalo Abecasis[48], Najaf Amin[49], Dan E Arking[50], Kristin L Ayers[51,52], Caterina M Barbieri[53], Chiara Batini[54], Joshua C Bis[55], Tineka Blake[54], Murielle Bochud[56], Michael Boehnke[25], Eric Boerwinkle[57], Dorret I Boomsma[58], Erwin P Bottinger[59], Peter S Braund[60,61], Marco Brumat[62], Archie Campbell[63,64], Harry Campbell[65], Aravinda Chakravarti[50], John C Chambers[1,5,66-68], Ganesh Chauhan[69], Marina Ciullo[70,71], Massimiliano Cocca[72], Francis Collins[73], Heather J Cordell[51], Gail Davies[74,75], Martin H de Borst[76], Eco J de Geus[58], Ian J Deary[74,75], Joris Deelen[77], Fabiola Del Greco M[78], Cumhur Yusuf Demirkale[79], Marcus Dörr[80,81], Georg B Ehret[50,82], Roberto Elosua[83,84], Stefan Enroth[85], A Mesut Erzurumluoglu[54], Teresa Ferreira[86,87], Mattias Frånberg[88-90], Oscar H Franco[91], Ilaria Gandin[62], Paolo Gasparini[62,72], Vilmantas Giedraitis[92], Christian Gieger[93-95], Giorgia Girotto[62,72], Anuj Goel[40,41], Alan J Gow[74,96], Vilmundur Gudnason[97,98], Xiuqing Guo[99], Ulf Gyllensten[85], Anders Hamsten[88,89], Tamara B Harris[100], Sarah E Harris[63,74], Catharina A Hartman[101], Aki S Havulinna[102,103], Andrew A Hicks[78], Edith Hofer[104,105], Albert Hofman[91,106], Jouke-Jan Hottenga[58], Jennifer E Huffman[19,107,108], Shih-Jen Hwang[107,108], Erik Ingelsson[109,110], Alan James[111,112], Rick Jansen[113], Marjo-Riitta Jarvelin[1,5,114-116], Roby Joehanes[107,117], Åsa Johansson[85], Andrew D Johnson[107,118], Peter K Joshi[65], Pekka Jousilahti[102], J Wouter Jukema[119],

Antti Jula[102], Mika Kähönen[120,121], Sekar Kathiresan[17,36,122], Bernard D Keavney[123,124], Kay-Tee Khaw[125], Paul Knekt[102], Joanne Knight[126], Ivana Kolcic[127], Jaspal S Kooner[5,67,68,128], Seppo Koskinen[102], Kati Kristiansson[102], Zoltan Kutalik[56,129], Maris Laan[130], Marty Larson[107], Lenore J Launer[100], Benjamin Lehne[1], Terho Lehtimäki[131,132], David CM Liewald[74,75], Li Lin[82], Lars Lind[133], Cecilia M Lindgren[40,87,134], YongMei Liu[135], Ruth JF Loos[28,59,136], Lorna M Lopez[74,137,138], Yingchang Lu[59], Leo-Pekka Lyytikäinen[131,132], Anubha Mahajan[40], Chrysovalanto Mamasoula[139], Jaume Marrugat[83], Jonathan Marten[19], Yuri Milaneschi[140], Anna Morgan[62], Andrew P Morris[40,141], Alanna C Morrison[142], Peter J Munson[79], Mike A Nalls[143,144], Priyanka Nandakumar[50], Christopher P Nelson[60,61], Teemu Niiranen[102,145], Ilja M Nolte[146], Teresa Nutile[70], Albertine J Oldehinkel[147], Ben A Oostra[49], Paul F O'Reilly[148], Elin Org[16], Sandosh Padmanabhan[64,149], Walter Palmas[150], Aarno Palotie[103,151,152], Alison Pattie[75], Brenda WJH Penninx[140], Markus Perola[102,103,153], Annette Peters[94,95,154], Ozren Polasek[127,155], Peter P Pramstaller[78,156,157], Quang Tri Nguyen[79], Olli T Raitakari[158,159], Rainer Rettig[161], Kenneth Rice[162], Paul M Ridker[23,163], Janina S Ried[94], Harriëtte Riese[147], Samuli Ripatti[103,164], Antonietta Robino[72], Lynda M Rose[23], Jerome I Rotter[99], Igor Rudan[165], Daniela Ruggiero[70,71], Yasaman Saba[166], Cinzia F Sala[53], Veikko Salomaa[102], Nilesh J Samani[60,61], Antti-Pekka Sarin[103], Reinhold Schmidt[104], Helena Schmidt[166], Nick Shrine[54], David Siscovick[167], Albert V Smith[97,98], Harold Snieder[146], Siim Sõber[130], Rossella Sorice[70], John M Starr[74,168], David J Stott[169], David P Strachan[170], Rona J Strawbridge[88,89], Johan Sundström[133], Morris A Swertz[171], Kent D Taylor[99], Alexander Teumer[81,172], Martin D Tobin[54], Maciej Tomaszewski[123,124], Daniela Toniolo[53], Michela Traglia[53], Stella Trompet[119,173], Jaakko Tuomilehto[174-177], Christophe Tzourio[21], André G Uitterlinden[91,178], Ahmad Vaez[146,179], Peter J van der Most[146], Cornelia M van Duijn[49], Germaine C Verwoert[91], Veronique Vitart[19], Uwe Völker[81,180], Peter Vollenweider[181], Dragana Vuckovic[62,182], Hugh Watkins[40,41], Sarah H Wild[183], Gonneke Willemsen[58], James F Wilson[19,65], Alan F Wright[19], Jie Yao[99], Tatijana Zemunik[184], Weihua Zhang[1,67], John R Attia[24], Adam S Butterworth[27,185], Daniel I Chasman[23,163], David Conen[186,187], Francesco Cucca[188,189], John Danesh[27,185], Caroline Hayward[19], Joanna MM Howson[27], Markku Laakso[190], Edward G Lakatta[191], Claudia Langenberg[28], Olle Melander[18], Dennis O Mook-Kanamori[26,192], Colin NA Palmer[32], Lorenz Risch[193-195], Robert A Scott[28], Rodney J Scott[24], Peter Sever[128], Tim D Spector[29], Pim van der Harst[196], Nicholas J Wareham[28], Eleftheria Zeggini[31], Daniel Levy[107,118], Patricia B Munroe[3,4], Christopher Newton-Cheh[134,197,198], Morris J Brown[3,4], Andres Metspalu[16], Bruce M. Psaty[201,202], Louise V Wain[54], Paul Elliott[1,5,203-205], Mark J Caulfield[3,4]

1. Department of Epidemiology and Biostatistics, Imperial College London, London, UK.
2. Department of Hygiene and Epidemiology, University of Ioannina Medical School, Ioannina, Greece.
3. William Harvey Research Institute, Barts and The London School of Medicine and Dentistry, Queen Mary University of London, London, UK.
4. National Institute for Health Research, Barts Cardiovascular Biomedical Research Center, Queen Mary University of London, London, UK.

5. MRC-PHE Centre for Environment and Health, Imperial College London, London, UK.
7. Division of Epidemiology, Department of Medicine, Institute for Medicine and Public Health, Vanderbilt Genetics Institute, Vanderbilt University Medical Center, Tennessee Valley Healthcare System (626)/Vanderbilt University, Nashville, TN, USA.
8. Vanderbilt Genetics Institute, Vanderbilt Epidemiology Center, Department of Obstetrics and Gynecology, Vanderbilt University Medical Center; Tennessee Valley Health Systems VA, Nashville, TN, USA.
9. Department of Epidemiology, Emory University Rollins School of Public Health, Atlanta, GA, USA.
10. Department of Biomedical Informatics, Emory University School of Medicine, Atlanta, GA, USA.
11. Massachusetts Veterans Epidemiology Research and Information Center (MAVERIC), VA Boston Healthcare System, Boston, USA.
12. Division of Aging, Department of Medicine, Brigham and Women's Hospital, Boston, MA, Department of Medicine, Harvard Medical School, Boston, MA, USA.
13. Atlanta VAMC and Emory Clinical Cardiovascular Research Institute, Atlanta, GA, USA.
14. VA Palo Alto Health Care System; Division of Cardiovascular Medicine, Stanford University School of Medicine, CA, USA.
15. Nephrology Section, Memphis VA Medical Center and University of Tennessee Health Science Center, Memphis, TN, USA.
16. Estonian Genome Center, University of Tartu, Tartu, Estonia.
17. Program in Medical and Population Genetics, Broad Institute of Harvard and MIT, Cambridge, MA, USA.
18. Department Clinical Sciences, Malmö, Lund University, Malmö, Sweden.
19. MRC Human Genetics Unit, MRC Institute of Genetics and Molecular Medicine, University of Edinburgh, Western General Hospital, Edinburgh, Scotland, UK
20. Department of Neurology, Bordeaux University Hospital, Bordeaux, France.
21. Univ. Bordeaux, Inserm, Bordeaux Population Health Research Center, CHU Bordeaux, Bordeaux, France.
22. Laboratory of Genetics and Genomics, NIA/NIH , Baltimore, MD, USA.
23. Division of Preventive Medicine, Brigham and Women's Hospital, Boston, MA, USA.
24. Hunter Medical Reseach Institute and Faculty of Health, University of Newcastle, New Lambton Heights, New South Wales, Australia.
25. Department of Biostatistics and Center for Statistical Genetics, University of Michigan, Ann Arbor, MI, USA.
26. Department of Clinical Epidemiology, Leiden University Medical Center, Leiden, the Netherlands.
27. MRC/BHF Cardiovascular Epidemiology Unit, Department of Public Health and Primary Care, University of Cambridge, Cambridge, UK.
28. MRC Epidemiology Unit, University of Cambridge School of Clinical Medicine, Cambridge, UK.
29. Department of Twin Research and Genetic Epidemiology, Kings College London, London, UK.

30. NIHR Biomedical Research Centre at Guy's and St Thomas' Foundation Trust, London, UK.
31. Wellcome Trust Sanger Institute, Hinxton, UK.
32. Division of Molecular and Clinical Medicine, School of Medicine, University of Dundee, UK.
33. Department of Pharmacy, COMSATS Institute of Information Technology, Abbottabad, Pakistan.
34. Department of Pathology and Molecular Medicine, McMaster University, Hamilton, Canada.
35. Institut universitaire de cardiologie et de pneumologie de Québec-Université Laval, , Quebec City, Canada.
36. Cardiovascular Research Center and Center for Human Genetic Research, Massachusetts General Hospital, Boston, Massachusetts, MA, USA.
37. University of Groningen, University Medical Center Groningen, Department of Cardiology, Groningen, The Netherlands.
38. University of Lille, Inserm, Centre Hosp. Univ Lille, Institut Pasteur de Lille, UMR1167 - RID-AGE - Risk factors and molecular determinants of aging-related diseases, Epidemiology and Public Health Department, Lille, France.
39. University of Dundee, Ninewells Hospital & Medical School, Dundee, , UK.
40. Wellcome Trust Centre for Human Genetics, University of Oxford, Oxford, UK.
41. Division of Cardiovascular Medicine, Radcliffe Department of Medicine, University of Oxford, Oxford, UK.
42. Usher Institute of Population Health Sciences and Informatics, University of Edinburgh, UK.
43. Department of Internal Medicine, Section Gerontology and Geriatrics, Leiden University Medical Center, Leiden, The Netherlands.
44. Imperial Clinical Trials Unit, Stadium House, 68 Wood Lane, London, UK.
45. School of Medicine, University College Dublin, Ireland.
46. Molecular and Cellular Therapeutics, Royal College of Surgeons in Ireland, Dublin, Ireland.
47. International Centre for Circulatory Health, Imperial College London, London, UK.
48. Center for Statistical Genetics, Dept. of Biostatistics, SPH II, Washington Heights, Ann Arbor, MI, USA.
49. Genetic Epidemiology Unit, Department of Epidemiology, Erasmus MC, Rotterdam, the Netherlands.
50. Center for Complex Disease Genomics, McKusick-Nathans Institute of Genetic Medicine, Johns Hopkins University School of Medicine, Baltimore, MD, USA.
51. Institute of Genetic Medicine, Newcastle University, Newcastle upon Tyne, UK.
52. Sema4, a Mount Sinai venture, Stamford, CT, USA.
53. Division of Genetics and Cell Biology, San Raffaele Scientific Institute, Milano, Italy.
54. Department of Health Sciences, University of Leicester, Leicester, UK.
55. Cardiovascular Health Research Unit, Department of Medicine, University of Washington, Seattle, WA, USA.
56. Institute of Social and Preventive Medicine, University Hospital of Lausanne, Lausanne, Switzerland.

57. Human Genetics Center, School of Public Health, The University of Texas Health Science Center at Houston and Human Genome Sequencing Center, Baylor College of Medicine, One Baylor Plaza, Houston, TX, USA.
58. Department of Biological Psychology, Vrije Universiteit Amsterdam, EMGO+ institute, VU University medical center, Amsterdam, the Netherlands.
59. The Charles Bronfman Institute for Personalized Medicine, Icachn School of Medicine at Mount Sinai, NY, USA.
60. Department of Cardiovascular Sciences, University of Leicester, Leicester, UK.
61. NIHR Leicester Biomedical Research Centre, Glenfield Hospital, Groby Road, Leicester, UK.
62. Department of Medical, Surgical and Health Sciences, University of Trieste, , Trieste, Italy.
63. Medical Genetics Section, Centre for Genomic and Experimental Medicine, Institute of Genetics and Molecular Medicine, University of Edinburgh, Edinburgh, UK.
64. Generation Scotland, Centre for Genomic and Experimental Medicine, University of Edinburgh, Edinburgh, UK.
65. Centre for Global Health Research, Usher Institute of Population Health Sciences and Informatics, University of Edinburgh, Edinburgh, Scotland, UK
66. Lee Kong Chian School of Medicine, Nanyang Technological University, Singapore, Singapore.
67. Department of Cardiology, Ealing Hospital, Middlesex, UK.
68. Imperial College Healthcare NHS Trust, London, UK.
69. Centre for Brain Research, Indian Institute of Science, Bangalore, India.
70. Institute of Genetics and Biophysics "A. Buzzati-Traverso", CNR, Napoli, Italy.
71. IRCCS Neuromed, Pozzilli, Isernia, Italy.
72. Institute for Maternal and Child Health IRCCS Burlo Garofolo, Trieste, Italy.
73. Medical Genomics and Metabolic Genetics Branch, National Human Genome Research Institute, NIH, Bethesda, MD, USA.
74. Centre for Cognitive Ageing and Cognitive Epidemiology, University of Edinburgh, 7 George Square, Edinburgh, UK.
75. Department of Psychology, University of Edinburgh, 7 George Square, Edinburgh, UK.
76. Department of Internal Medicine, Division of Nephrology, University of Groningen, University Medical Center Groningen, Groningen, The Netherlands.
77. Department of Molecular Epidemiology, Leiden University Medical Center, Leiden, the Netherlands.
78. Institute for Biomedicine, Eurac Research, Bolzano, Italy - Affiliated Institute of the University of Lübeck, Lübeck, Germany.
79. Mathematical and Statistical Computing Laboratory, Office of Intramural Research, Center for Information Technology, National Institutes of Health, Bethesda, MD, USA.
80. Department of Internal Medicine B, University Medicine Greifswald, Greifswald, Germany.
81. DZHK (German Centre for Cardiovascular Research), partner site Greifswald, Greifswald, Germany.
82. Cardiology, Department of Medicine, Geneva University Hospital, Geneva, Switzerland.

83. CIBERCV & Cardiovascular Epidemiology and Genetics, IMIM. Dr Aiguader 88, Barcelona, Spain.
84. Faculty of Medicine, Universitat de Vic-Central de Catalunya, Vic, Spain.
85. Department of Immunology, Genetics and Pathology, Uppsala Universitet, Science for Life Laboratory, Uppsala, Sweden.
86. Wellcome Centre for Human Genetics, University of Oxford, Roosevelt Drive, Oxford, UK.
87. Big Data Institute, Li Ka Shing Center for Health for Health Information and Discovery, Oxford University, Old Road, Oxford, UK.
88. Cardiovascular Medicine Unit, Department of Medicine Solna, Karolinska Institutet, Stockholm, Sweden.
89. Centre for Molecular Medicine, L8:03, Karolinska Universitetsjukhuset, Solna, Sweden.
90. Department of Numerical Analysis and Computer Science, Stockholm University, Stockholm, Sweden.
91. Department of Epidemiology, Erasmus MC, Rotterdam, the Netherlands.
92. Department of Public Health and Caring Sciences, Geriatrics, Uppsala, Sweden.
93. Research Unit of Molecular Epidemiology, Helmholtz Zentrum München, German Research Center for Environmental Health, Neuherberg, Germany.
94. Institute of Epidemiology, Helmholtz Zentrum München, German Research Center for Environmental Health, Neuherberg, Germany.
95. German Center for Diabetes Research (DZD e.V.), Neuherberg, Germany.
96. Department of Psychology, School of Social Sciences, Heriot-Watt University, Edinburgh, UK.
97. Faculty of Medicine, University of Iceland, Reykjavik, Iceland.
98. Icelandic Heart Association, Kopavogur, Iceland.
99. The Institute for Translational Genomics and Population Sciences, Department of Pediatrics, LABioMed at Harbor-UCLA Medical Center, Torrance, CA, USA.
100. Intramural Research Program, Laboratory of Epidemiology, Demography, and Biometry, National Institute on Aging, Bethesda, MD, USA.
101. Department of Psychiatry, University of Groningen, University Medical Center Groningen, Groningen, The Netherlands.
102. Department of Public Health Solutions, National Institute for Health and Welfare (THL), Helsinki, Finland.
103. Institute for Molecular Medicine Finland (FIMM), University of Helsinki, Helsinki, Finland.
104. Clinical Division of Neurogeriatrics, Department of Neurology, Medical University of Graz, Graz, Austria.
105. Institute for  Medical Informatics, Statistics and Documentation, Medical University of Graz, Graz, Austria.
106. Department of Epidemiology, Harvard T.H. Chan School of Public Health, Boston, MA, USA.
107. National Heart, Lung and Blood Institute's Framingham Heart Study, Framingham, MA, USA.
108. The Population Science Branch, Division of Intramural Research, National Heart Lung and Blood Institute national Institute of Health, Bethesda, MD, USA.

109. Department of Medical Sciences, Molecular Epidemiology and Science for Life Laboratory, Uppsala University, Uppsala, Sweden.
110. Division of Cardiovascular Medicine, Department of Medicine, Stanford University School of Medicine, Stanford, CA USA.
111. Department of Pulmonary Physiology and Sleep, Sir Charles Gairdner Hospital, Hospital Avenue, Nedlands, Australia.
112. School of Medicine and Pharmacology, University of Western Australia.
113. Department of Psychiatry, VU University Medical Center, Amsterdam Neuroscience, Amsterdam, the Netherlands.
114. Biocenter Oulu, University of Oulu, Oulu, Finland.
115. Center For Life-course Health Research, University of Oulu, Oulu Finland.
116. Unit of Primary Care, Oulu University Hospital, Oulu, Oulu, Finland.
117. Hebrew SeniorLife, Harvard Medical School, Boston, MA, USA.
118. Population Sciences Branch, National Heart, Lung and Blood Institute, National Institutes of Health, Bethesda, MD, USA.
119. Department of Cardiology, Leiden University Medical Center, Leiden, the Netherlands.
120. Department of Clinical Physiology, Tampere University Hospital, Tampere, Finland.
121. Department of Clinical Physiology, Finnish Cardiovascular Research Center - Tampere, Faculty of Medicine and Life Sciences, University of Tampere, Tampere, Finland.
122. Broad Institute of the Massachusetts Institute of Technology and Harvard University, Cambridge, MA, USA.
123. Division of Cardiovascular Sciences, Faculty of Biology, Medicine and Health, The University of Manchester, Manchester, UK.
124. Division of Medicine, Manchester University NHS Foundation Trust, Manchester Academic Health Science Centre, Manchester, UK
125. Department of Public Health and Primary Care, Institute of Public Health, University of Cambridge, Cambridge, UK.
126. Data Science Institute and Lancaster Medical School, Lancaster, UK.
127. Department of Public Health, Faculty of Medicine, University of Split, Croatia.
128. National Heart and Lung Institute, Imperial College London, London, UK.
129. Swiss Institute of Bioinformatics, Lausanne, Switzerland.
130. Institute of Biomedicine and Translational Medicine, University of Tartu, Tartu, Estonia.
131. Department of Clinical Chemistry, Fimlab Laboratories, Tampere, Finland.
132. Department of Clinical Chemistry, Finnish Cardiovascular Research Center - Tampere, Faculty of Medicine and Life Sciences, University of Tampere, Tampere, Finland
133. Department of Medical Sciences, Cardiovascular Epidemiology, Uppsala University, Uppsala, Sweden.
134. Program in Medical and Population Genetics, Broad Institute, Cambridge, MA, USA.
135. Division of Public Health Sciences, Wake Forest School of Medicine, Winston-Salem, NC, USA.
136. Mindich Child health Development Institute, The Icahn School of Medicine at Mount Sinai, New York, NY, USA.
137. Department of Psychiatry, Royal College of Surgeons in Ireland, Education and Research Centre, Beaumont Hospital, Dublin, Ireland.

138. University College Dublin, UCD Conway Institute, Centre for Proteome Research, UCD, Belfield, Dublin, Ireland.
139. Institute of Health and Society, Newcastle University, Newcastle upon Tyne, UK.
140. Department of Psychiatry, Amsterdam Public Health and Amsterdam Neuroscience, VU University Medical Center/GGZ inGeest, Amsterdam, The Netherlands.
141. Department of Biostatistics, University of Liverpool, Block F, Waterhouse Building, Liverpool, UK.
142. Department of Epidemiology, Human Genetics and Environmental Sciences, School of Public Health, University of Texas Health Science Center at Houston, Houston, TX, USA.
143. Data Tecnica International, Glen Echo, MD, USA.
144. Laboratory of Neurogenetics, National Institute on Aging, Bethesda, USA.
145. Department of Medicine, Turku University Hospital and University of Turku, Finland.
146. Department of Epidemiology, University of Groningen, University Medical Center Groningen, Groningen, The Netherlands.
147. Interdisciplinary Center Psychopathology and Emotion regulation (ICPE), University of Groningen, University Medical Center Groningen, Groningen, The Netherlands.
148. SGDP Centre, Institute of Psychiatry, Psychology and Neuroscience, King's College London, London, UK.
149. British Heart Foundation Glasgow Cardiovascular Research Centre, Institute of Cardiovascular and Medical Sciences, College of Medical, Veterinary and Life Sciences, University of Glasgow, Glasgow, UK.
150. Department of Medicine, Columbia University Medical Center, New York, NY, USA.
151. Analytic and Translational Genetics Unit, Department of Medicine, Department of Neurology and Department of Psychiatry Massachusetts General Hospital, Boston, MA, USA.
152. The Stanley Center for Psychiatric Research and Program in Medical and Population Genetics, The Broad Institute of MIT and Harvard, Cambridge, MA, USA.
153. University of Tartu, Tartu, Estonia.
154. German Center for Cardiovascular Disease Research (DZHK), partner site Munich, Neuherberg, Germany.
155. Psychiatric hospital “Sveti Ivan”, Zagreb, Croatia.
156. Department of Neurology, General Central Hospital, Bolzano, Italy.
157. Department of Neurology, University of Lübeck, Lübeck, Germany.
158. Department of Clinical Physiology and Nuclear Medicine, Turku University Hospital, Turku, Finland.
159. Research Centre of Applied and Preventive Cardiovascular Medicine, University of Turku, Turku, Finland.
161. Institute of Physiology, University Medicine Greifswald, Karlsburg, Germany.
162. Department of Biostatistics University of Washington, Seattle, WA, USA.
163. Harvard Medical School, Boston MA.
164. Public health, Faculty of Medicine, University of Helsinki, Finland
165. Centre for Global Health Research, Usher Institute of Population Health Sciences and Informatics, University of Edinburgh, Scotland, UK.

166. Gottfried Schatz Research Center for Cell Signaling, Metabolism & Aging, Molecular Biology and Biochemistry, Medical University of Graz, Graz, Austria.
167. The New York Academy of Medicine, New York, NY, USA.
168. Alzheimer Scotland Dementia Research Centre, University of Edinburgh, Edinburgh, UK.
169. Institute of Cardiovascular and Medical Sciences, Faculty of Medicine, University of Glasgow, United Kingdom.
170. Population Health Research Institute, St George's, University of London, London, UK.
171. Department of Genetics, University of Groningen, University Medical Center Groningen, Groningen, The Netherlands.
172. Institute for Community Medicine, University Medicine Greifswald, Greifswald, Germany.
173. Department of Gerontology and Geriatrics, Leiden University Medical Center, Leiden, the Netherlands.
174. Dasman Diabetes Institute, Dasman, Kuwait.
175. Chronic Disease Prevention Unit, National Institute for Health and Welfare, Helsinki, Finland.
176. Department of Public Health, University of Helsinki, Helsinki, Finland.
177. Saudi Diabetes Research Group, King Abdulaziz University, Jeddah, Saudi Arabia.
178. Department of Internal Medicine, Erasmus MC, Rotterdam, the Netherlands.
179. Research Institute for Primordial Prevention of Non-communicable Disease, Isfahan University of Medical Sciences, Isfahan, Iran.
180. Interfaculty Institute for Genetics and Functional Genomics, University Medicine Greifswald, Greifswald, Germany.
181. Department of Internal Medicine, University Hospital, CHUV, Lausanne, Switzerland.
182. Experimental Genetics Division, Sidra Medical and Research Center, Doha, Qatar.
183. Centre for Population Health Sciences, Usher Institute of Population Health Sciences and Informatics, University of Edinburgh, Scotland, UK
184. Department of Biology, Faculty of Medicine, University of Split, Croatia.
185. The National Institute for Health Research Blood and Transplant Research Unit in Donor Health and Genomics, University of Cambridge, UK.
186. Division of Cardiology, University Hospital, Basel, Switzerland.
187. Division of Cardiology, Department of Medicine, McMaster University, Hamilton, Canada.
188. Institute of Genetic and Biomedical Research, National Research Council (CNR), Monserrato, Cagliari, Italy.
189. Department of Biomedical Sciences, University of Sassari, Sassari, Italy.
190. Institute of Clinical Medicine, Internal Medicine, University of Eastern Finland and Kuopio University Hospital, Kuopio, Finland.
191. Laboratory of Cardiovascular Science, NIA/NIH , Baltimore, MD, USA.
192. Department of Public Health and Primary Care, Leiden University Medical Center, Leiden, the Netherlands.
193. Labormedizinisches Zentrum Dr. Risch, Schaan, Liechtenstein.
194. Private University of the Principality of Liechtenstin, Triesen, Liechtenstein.

195. University Insitute of Clinical Chemistry, Inselspital, Bern University Hospital, University of Bern, Bern, Switzerland.
196. Department of Cardiology, University of Groningen, University Medical Center Groningen, Groningen, The Netherlands.
197. Center for Genomic Medicine, Massachusetts General Hospital, Boston, MA, USA.
198. Cardiovascular Research Center, Massachusetts General Hospital, Boston, MA, USA.
201. Cardiovascular Health Research Unit, Departments of Medicine, Epidemiology and Health Services, University of Washington, Seattle, WA, USA.
202. Kaiser Permanente Washington Health Research Institute, Seattle, WA, USA.
203. National Institute for Health Research Imperial Biomedical Research Centre, Imperial College Healthcare NHS Trust and Imperial College London, London, UK.
204. UK Dementia Research Institute (UK DRI) at Imperial College London, London, UK
205. Health Data Research-UK London substantive site, London, U.K
